# PD-1 in hippocampal neurons regulates excitability, synaptic plasticity, and cognition

**DOI:** 10.1101/2021.07.13.451894

**Authors:** Junli Zhao, Sangsu Bang, Aidan McGinnis, Kenta Furutani, Changyu Jiang, Alexus Roberts, Christopher R Donnelly, Qianru He, Mei-Chuan Ko, Haichen Wang, Richard D. Palmiter, Ru-Rong Ji

**Affiliations:** Center for Translational Pain Medicine, Department of Anesthesiology, Duke University Medical Center, Durham, North Carolina, USA; Department of Physiology and Pharmacology, Wake Forest School of Medicine, Winston-Salem, NC, USA; Department of Neurology, Duke University Medical Center, Durham, North Carolina, USA; Department of Biochemistry, University of Washington, Seattle, Washington, USA; Howard Hughes Medical Institute, University of Washington, Seattle, WA, USA; Department of Cell Biology, Duke University Medical Center, Durham, North Carolina, USA; Department of Neurobiology, Duke University Medical Center, Durham, North Carolina, USA

**Keywords:** programmed cell death protein 1, hippocampal neurons, immunotherapy, long-term potentiation, microglia, mice, nonhuman primate, traumatic brain injury

## Abstract

Immunotherapy using monoclonal antibodies against programmed cell death protein 1 (PD-1) demonstrated improved survival in cancer patients through immune activation. Here we show that functional PD-1 is expressed in mouse and primate hippocampal neurons and PD-1 inhibition improves cognition in physiological and pathological conditions. Mice lacking the *Pdcd1* gene encoding PD-1 exhibit enhanced long-term potentiation (LTP) and learning and memory. These behavioral and cellular changes can be recapitulated by selective deletion of *Pdcd1* in hippocampal excitatory neurons but not in microglia. Perfusion of mouse or nonhuman primate brain slices with anti-PD-1 antibody is sufficient to increase excitability in CA1 hippocampal neurons. Conversely, re-expression of *Pdcd1* in PD-1 deficient hippocampal neurons suppresses memory and LTP. Traumatic brain injury impairs learning and memory, which is improved by intraventricular administration of anti-PD-1. These findings suggest that anti-PD-1 treatment has therapeutic potential to counteract cognitive decline.

**Highlights:**

- Adult mice lacking *Pdcd1* in hippocampal neurons exhibit enhanced memory and LTP
- Anti-PD-1 antibody treatment increases CA1 neuron excitability in brain slices of mice and primates
- Re-expression of *Pdcd1* in PD-1 deficient hippocampal neurons impairs memory and LTP
- Cognitive deficits after traumatic brain injury are improved by anti-PD-1 treatment

## INTRODUCTION

Programmed cell death protein 1 (PD-1), also known as CD279, is a surface molecule expressed by immune cells such as T cells. As an immune checkpoint regulator PD-1 down-regulates the immune system and suppresses T cell activities, and therefore, serves as a primary target of immunotherapy agents that fight cancer (Ishida et al., 1992; Sharma and Allison, 2015). New evidence suggests that PD-1 also has an active role in the nervous system. For example, systemic PD-1 blockade promoted clearance of cerebral amyloid-β plaques and improved memory in mouse models of Alzheimer’s disease (AD) (Baruch et al., 2016; Rosenzweig et al., 2019) by recruitment of peripheral macrophages into the central nervous system (CNS). However, other studies reported that systemic anti-PD-1 treatment failed to affect Aβ pathology in three different models of AD (Latta-Mahieu et al., 2018) and PD-1 deficiency did not induce myeloid mobilization to the brain during chronic neurodegeneration (Obst et al., 2018). It was also reported that chronic PD-1 checkpoint blockade (weekly treatment for 12 weeks) did not affect cognition or promote tau clearance in a tauopathy mouse model, although it increased locomotor activity (Lin et al., 2019). Another study showed that immunotherapy using influenza vaccine combined with a moderate-dose PD-1 blockade reduced amyloid-β accumulation and improved cognition in mouse AD model through recruitment of peripheral monocyte-derived macrophages into the brain (Xing et al., 2021). Notably, these contradictory studies exclusively focused on immunoregulation of PD-1; direct regulation of neuronal PD-1 was not addressed.

We recently demonstrated that primary sensory neurons in the peripheral nervous system (PNS), as well as spinal cord neurons in the CNS, express *Pdcd1* mRNA and functional PD-1 protein (Chen et al., 2017; Jiang et al., 2020; Wang et al., 2020b). Notably, PD-1 functions to suppress neuronal excitability, as *Pdcd1-*deficient mice show hyperexcitability in nociceptive sensory neurons, leading to pain hypersensitivity (Chen et al., 2017) and impaired morphine analgesia in mice (Wang et al., 2020b). Neuronal expression of PD-1 is also present in the brain. For example, single-cell analysis revealed low expression levels of *Pdcd1* mRNA in excitatory cortical neurons (Zeisel et al., 2018). We found that *Pdcd1* mRNA and PD-1 protein are widely expressed by neurons in different brain regions included cortex, thalamus, and hypothalamus; furthermore, PD-1 is required for GABA receptor signaling in these brain regions and acts as a neuronal activity inhibitor in the CNS to regulate the balance of excitation and inhibition under the physiological conditions (Jiang et al., 2020). PD-1 activation could suppress the phosphorylation of extracellular signal-regulated kinase (ERK) in neurons, a critical process in synaptic plasticity underlying learning and memory (Araki et al., 2015; Atkins et al., 1998; Impey et al., 1998). However, we could not exclude a possible involvement of glial cells in these previous studies, as microglia are also known to express PD-1 (Suda et al., 2021; Yao et al., 2014) and regulate synaptic plasticity and memory (Parkhurst et al., 2013; Salter and Stevens, 2017). To determine the specific role of PD-1 in neurons, we removed PD-1 function selectively in neurons or microglia by generating *Pdcd1* conditional knockout (cKO) mice. Using these mice, we show that PD-1 also plays a critical role in regulating synaptic function in hippocampal neurons and regulates learning and memory. We also demonstrate that infusion of antibodies against PD-1 improves cognition after traumatic brain injury (TBI); hence, this approach has potential as a neurotherapy.

## RESULTS

### *Pdcd1* deficiency or PD-1 blockade enhances learning and memory in mice

To determine whether PD-1 regulates learning and memory, we first used the novel object recognition (NOR) test to compare the discrimination index in wild-type (WT) and global knockout mice lacking *Pdcd1* (PD-1 KO, Figure S1A). Compared with age-matched WT mice (8 to 12 weeks old), PD-1 KO mice spent more time exploring the novel object and exhibited a higher discrimination index at 0.5 h and 24 h (Figures 1A, B), indicating increased learning and memory in this test. We also examined spatial learning and memory in WT and PD-1 KO mice using the Morris Water maze (MWM) testing (Figure S1B). During hidden platform training, PD-1 KO mice spent less time navigating to the platform than WT mice (Figure 1C), suggesting enhanced spatial learning ability. In the MWM probe test, the latency to find the platform was substantially reduced in PD-1 KO mice (WT: 29.41 ± 5.127 s vs. PD-1 KO: 8.480 ± 2.138 s), whereas the number of crossing was markedly increased in KO mice (Figures 1D and 1E); these results support an improvement in spatial memory by PD-1 KO mice. Notably, this deficiency did not alter the motor function in the rotarod test, anxiety in the open-field or elevate-plus maze tests, but PD-1 KO mice were less depressed in the tail suspension test (Figures S1J-S1N).

**Figure 1.**
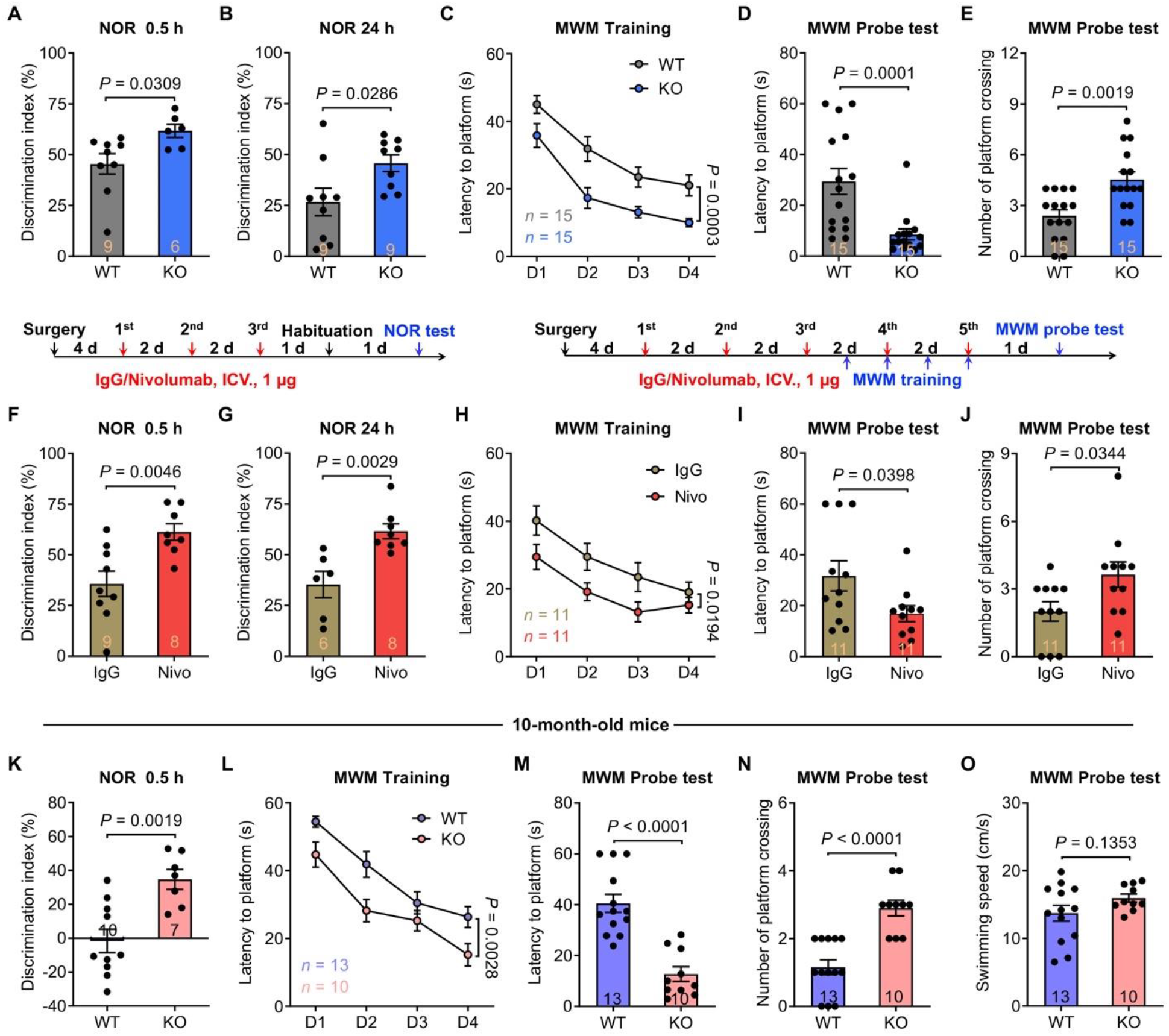
PD-1 loss or blockade enhances learning and memory. **(A-B)** Novel object recognition (NOR) test in WT mice (n = 9) and PD-1 KO mice (n = 6 and 9) at 0.5 h (A) and 24 h (B). Note that PD-1 KO mice exhibit higher discrimination index in both short-term (0.5 h, *P* = 0.0309) and long-term (24 h, *P* = 0.0286) tests. (**C)** Spatial learning curves during Morris water maze (MWM) training in WT mice (n =15) and PD-1 KO mice (n = 15), measured as latency to find the hidden platform. Note that PD-1 KO mice spend less time locating the hidden platform compared to WT mice (*P* = 0.0003). **(D-E)** MWM probe test for latency to platform (D) and number of platform crossings (E) in WT mice (n = 15) and PD-1 KO mice (n = 15). PD-1 KO mice have shorter latency (D, *P* = 0.0001) and more platform zone crossings (E, *P* = 0.0019) than WT mice. **(F-G)** NOR test in WT mice treated with control IgG (n = 9 and 6) and nivolumab (n = 8) at 0.5 h (F) and 24 h (G). Nivolumab treatment significantly increases discrimination index in short-term (0.5 h, *P* = 0.0046) and long-term (24 h, *P* = 0.0029) tests. Control IgG and nivolumab (3 × 1 μg) was ICV administrated every other day. (**H**) Spatial learning curves during MWM training period show significant improvement (*P* = 0.0194) in WT mice treated with nivolumab than control IgG (n = 11). **(I-J)** MWM probe test for latency to platform (I) and number of platform crossings (J) in WT mice with control IgG and nivolumab (n = 11). Nivolumab significantly improves the platform locating time (*P* = 0.0398) and the platform zone crossings number (*P* = 0.0344). Control IgG and nivolumab (5 × 1 μg) was ICV administrated every other day (H-J). **(K-O)** Distinct cognitive function in 10-month-old WT and PD-1 KO mice. **(K)** NOR test in WT mice (n = 10) and PD-1 KO mice (n = 7). Dementia is improved in PD-1 KO mice (*P* = 0.0019). **(L)** During MWM training phase PD-1 KO mice (n = 10) show stronger ability to locate the hidden platform than WT mice (n = 13, *P* = 0.0028). **(M-N)** MWM probe test shows greater ability of PD-1 KO mice (n = 10) vs. WT mice (n = 13) in locating the platform zone (M, *P* < 0.0001) and crossing the platform zone (N, *P* < 0.0001). **(O)** In MWM probe test WT (n = 13) and PD-1 KO (n = 10) mice are comparable in swimming speed (*P* = 0.1353). Data are represented as mean ± SEM. Two-tailed Student’s t test (A, B, F, G, K, O), Two-way ANOVA (C, H, L), Mann-Whitney test (D, E, I, J, M, N). Also see **Figure S1**.

To circumvent the possibility of developmental compensation in PD-1 KO mice, we examined whether central PD-1 blockade in WT adult mice could recapitulate the phenotypes observed in PD-1 KO mice. Nivolumab is an FDA-approved fully humanized IgG monoclonal antibody that binds to mouse PD-1 and produces a functional blockade in behavioral assays (Chen et al., 2017). We administrated nivolumab at a very low dose (1 µg ≈ 7 pmol, 3 times, every other day) to WT mice by intraventricular (ICV) injection, thereby targeting PD-1 function in the brain. This treatment increased the discrimination index in NOR tests at 0.5 h and 24 h (Figures 1F and 1G), as compared to treatment with human IgG as control. We also examined spatial learning and memory after ICV injections (5 × 1 µg, every other day) of nivolumab. In hidden the platform training phase, nivolumab-treated WT mice exhibited a shorter latency to find the platform than control IgG-injected mice (Figure 1H). In the probe test, nivolumab-treated WT mice had better performance in platform location navigation (Figure 1I) and an increased number of platform crossings (Figure 1J). Swimming speed was slightly increased after *Pdcd1* deletion but not after nivolumab treatment (Figures S1C and S1D), suggesting that learning and memory improvement is not a result of enhanced swimming capability. Notably, nivolumab-induced learning and memory improvement was abolished in PD-1 KO mice (Figures S1E-S1I), confirming a specific effect of nivolumab on PD-1. Together, these data indicate that either genetic deletion of *Pdcd1* in global KO mice or PD-1 blockade with nivolumab in WT adult mice is sufficient to enhance learning and memory, highlighting PD-1 as a negative regulator of cognition.

We also investigated learning and memory in 10-month-old WT and PD-1 KO mice with cognitive decline. These older WT mice had a mean discrimination index of −1.588 ± 6.916 (Figure 1K), a sign of cognitive decline compared to young mice. However, the age-matched PD-1 KO mice had a higher discrimination index (mean = 34.66 ± 6.83) in this test (Figure 1K). The 10-month-old PD-1 KO mice also had better performance in the MWM tests compared with age-matched WT mice (Figures 1L-1N), with no difference in swimming speed (Figure 1O). These data suggest that PD-1 contributes to dementia in 10-month-old mice.

### Loss of PD-1 increases neuronal excitability in hippocampal neurons

The hippocampus plays a crucial role in learning and memory (Malenka et al., 1989; Volk et al., 2013). Single-cell RNA-seq revealed low-levels of *Pdcd1* mRNA in cortical and hippocampal neurons (Zeisel et al., 2018). Using sensitive RNAscope *in situ* hybridization, we demonstrated widespread expression of *Pdcd1* mRNA in cortical and thalamic neurons (Jiang et al., 2020). We also examined the expression of PD-1 protein and *Pdcd1* mRNA in the hippocampus. Double immunostaining revealed PD-1 expression by 20-40% hippocampal CA1 and CA3 neurons that co-express neuronal maker NeuN (Figure 2A, Figures S2A and S2C). Notably, this PD-1 immunostaining was abolished in the presence of a blocking peptide as well as in PD-1 KO mice (Figure 2A and Figure S2A). *In situ* hybridization showed *Pdcd1* expression in 55-65% CA1 and CA3 neurons and this expression was absent in PD-1 KO mice (Figure 2B and Figures S2B and S2D).

**Figure 2.**
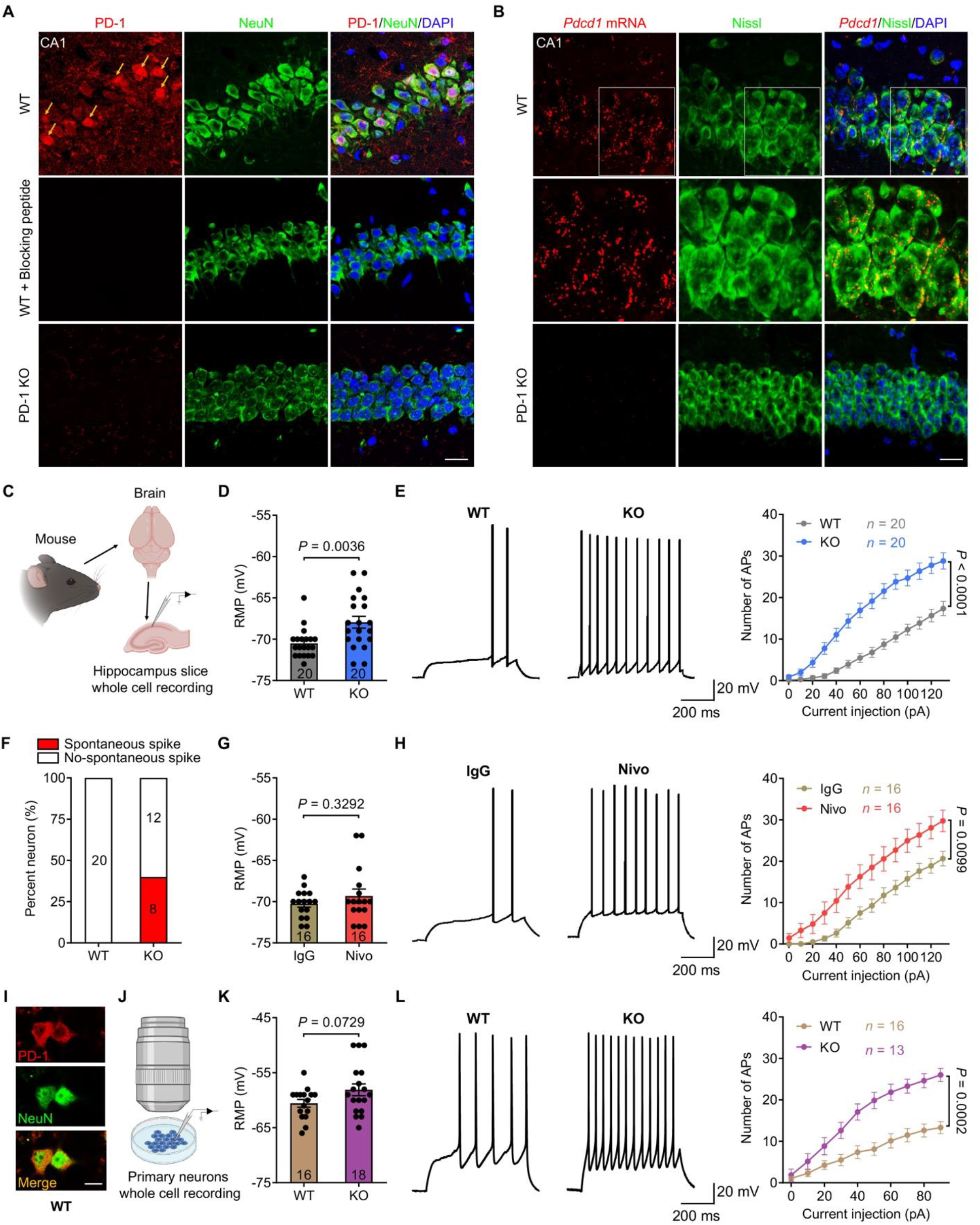
PD-1 is expressed by mouse hippocampal neurons and PD-1 loss or blockade increases neuronal excitability in hippocampal neurons. **(A)** Representative images of immunostaining show expression of PD-1 in WT neurons, which is absent after co-incubation with a PD-1 blocking peptide or in PD-1 KO CA1 neurons. Arrows indicate PD-1 positive neurons. Scale bar, 20 µm. **(B)** Representative images of *in situ* hybridization show expression of *Pdcd1* in WT but not in PD-1 KO CA1 neurons. Higher magnification images (middle) show *Pdcd1* in Nissl-labeled WT CA1 neurons. Scale bar, 20 µm. **(C)** Schematic of experimental design for whole-cell patch clamp recordings in mouse CA1 neurons. **(D)** RMP recorded in WT (n = 20 neurons from 6 mice) and PD-1 KO (n = 20 neurons from 3 mice) CA1 neurons. Note that depolarized RMP occurred in PD-1 KO CA1 neurons (*P* = 0.0036). **(E)** Representative AP traces and the number of APs evoked by step current injection in WT (n = 20 neurons from 6 mice) and PD-1 KO (n = 20 neurons from 3 mice) CA1 neurons. Neuronal excitability is increased in PD-1 KO CA1 neurons (*P* < 0.0001). **(F)** Spontaneous discharges present in PD-1 KO CA1 neurons (8/20) but not in WT CA1 neurons (0/20). **(G)** RMP recorded in WT CA1 neurons with incubation of control IgG (n = 16 neurons from 3 mice) or nivolumab (n = 20 neurons from 4 mice). RMP is comparable in these two groups (*P* = 0.3292). **(H)** Representative AP traces and the number of APs evoked by step current injection in CA1 neurons treated with control IgG (n = 16 neurons from 3 mice) or nivolumab (n = 20 neurons from 4 mice). Nivolumab-treated CA1 neurons show increased neuronal excitability than control IgG (*P* = 0.0099). **(I)** Representative images of PD-1 immunocytochemistry in WT primary hippocampal neurons. Scale bar, 10 µm. **(J)** Schematic depicting patch clamp recordings in cultured primary hippocampal neurons from WT and PD-1 KO mice. **(K)** RMP recorded in WT (n = 16 neurons from 3 mice/18 embryos) and PD-1 KO (n = 18 neurons from 2 mice/10 embryos) primary hippocampal neurons. Note that RMP of PD-1 KO neurons is moderately increased (*P* = 0.0729). **(L)** Representative AP traces and the number of APs evoked by step current injection in WT (n = 16 neurons from 3 mice/18 embryos) and PD-1 KO (n = 13 neurons from 2 mice/10 embryos) primary hippocampal neurons. Neuronal excitability is increased in PD-1 KO neurons (*P* = 0.0002). Data are represented as mean ± SEM. Two-tailed Student’s t test (D, G, K), Two-way ANOVA (E, H, L). Also see **Figure S2**.

To define the function of PD-1 in hippocampal neurons, we first examined the neuronal excitability in CA1 neurons of WT and PD-1 KO mice using *ex vivo* whole-cell, current-clamp patch recording in brain slices (Figure 2C). Compared to WT neurons, CA1 pyramidal neurons from PD-1 KO mice had increased resting membrane potential (RMP), suggesting a cell membrane depolarization (Figure 2D). Furthermore, PD-1 KO CA1 neurons fired more action potentials (APs) evoked by current injection (0-130 pA, 10 pA step, Figure 2E). Intriguingly, we observed spontaneous discharges in 40% of KO CA1 neurons (8/20) but none in WT neurons (0/20, Figure 2F). Additionally, anti-PD-1 treatment of WT slices also increased neuronal excitability. Compared with control IgG, acute incubation of brain slices with nivolumab (300 ng/mL = 7 nM, 2 h) did not change the RMP but significantly increased AP firing rate in CA1 neurons (Figures 2G and 2H). Thus, loss of PD-1 results in increased neuronal excitability in CA1 neurons.

To further determine the role of PD-1 in CA1 neurons, we dissociated hippocampal neurons in primary cultures from E17-E19 mice (Figures 2I-2L). Immunocytochemistry in cultured primary hippocampal neurons revealed PD-1 expression in NeuN^+^ neurons (Figure 2I) and nivolumab did not bind to PD-1 KO hippocampal neurons (Figure S2E). Patch-clamp recording in dissociated hippocampal neurons (7-8 days in vitro, DIV) showed that PD-1 KO neurons had a moderate increase in RMP (Figure 2K) but a marked increase in AP firing rate following current injection compared to WT cultures (Figure 2L). Thus, PD-1 on CA1 neurons normally dampens neuronal excitability.

### Loss of PD-1 increases hippocampal synaptic function and dendritic spine density

We used additional electrophysiological and histochemical approaches to study how PD-1 depletion affects hippocampal synaptic functions. Whole-cell, voltage-patch-clamp recording in brain slices revealed increases in the frequency and amplitude of spontaneous miniature excitatory postsynaptic current (mEPSC) in PD-1 KO CA1 neurons compared to WT neurons (Figure 3A). The increase in mEPSC frequency was recapitulated by PD-1 blockade with nivolumab (300 ng/mL, 2 h) in WT slices (Figure 3B).

**Figure 3.**
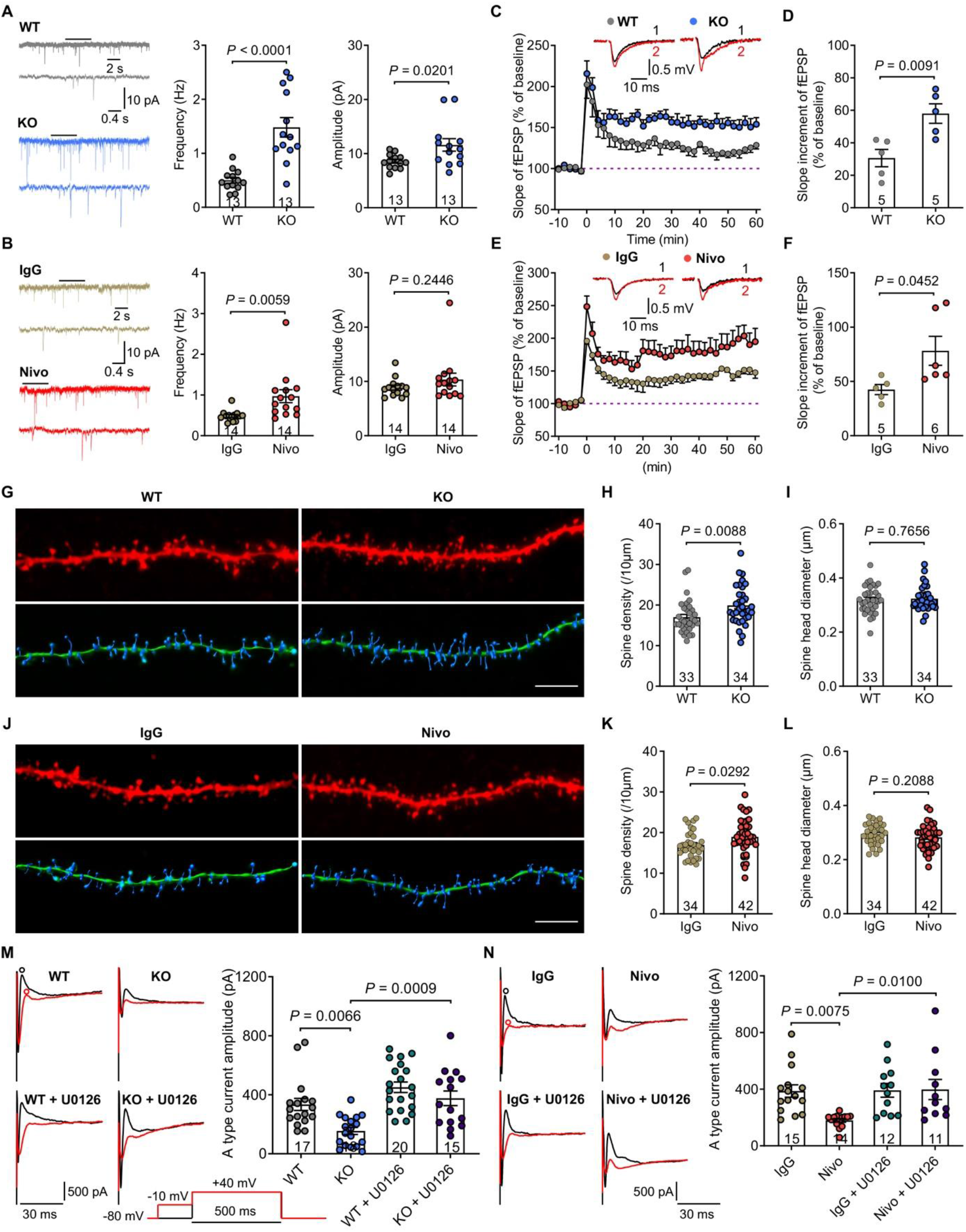
PD-1 deficiency increases hippocampal synaptic transmission, synaptic plasticity and dendritic spine density, and reduces A type potassium currents. **(A)** Representative traces (left) and quantification of mEPSC frequency (middle, *P* < 0.0001) and amplitude (right, *P* = 0.0201) of in WT (n = 13 neurons from 3 mice) and PD-1 KO (n = 13 neurons from 4 mice) CA1 neurons. **(B)** Representative traces (left) and quantification of frequency (middle, *P* = 0.0059) and amplitude (right, *P* = 0.2446) of mEPSC in control IgG (n = 14 neurons from 4 mice) and nivolumab (n = 14 neurons from 3 mice) treated CA1 neurons. Low panel traces in A and B are enlargements of the top panel traces indicted by bars. (**C-D**) Summary plots (C) and average slope (D) of LTP induced by 2 × HFS in the CA1 region of WT slices (n = 5 from 5 mice) and PD-1 KO slices (n = 5 from 5 mice). PD-1 KO slices show enhanced slope of fEPSP than WT slices (*P* = 0.0091). Top: Black traces (1) and red trace (2) represent the baseline fEPSP and post-induction fEPSP, respectively. **(E-F)** Summary plots (E) and average slope (F) of LTP induced by 2 × HFS in the CA1 region of control IgG-treated slices (n = 5 from 5 mice) and nivolumab-treated slices (n = 6 from 5 mice). Note that nivolumab treatment enlarges the slope of fEPSP (*P* = 0.0452). **(G)** Representative confocal z stack and three-dimensional reconstruction images of the apical dendrites of CA1 neurons obtained from WT and PD-1 KO slices. Scale bar, 5 μm. **(H-I)** Statistical analyses of spine density (H, *P* = 0.0088) and spine size (I, *P* = 0.7656) in CA1 neurons from WT mice (n = 33 dendrites from 4 mice) and PD-1 KO (n = 34 dendrites from 4 mice). **(J)** Representative confocal z stack and three-dimensional reconstruction images of the apical dendrites of CA1 neurons obtained from brains treated with control IgG and nivolumab. Scale bar, 5 μm. **(K-L)** Statistical analyses of spine density (K, *P* = 0.0292) and spine size (L, *P* = 0.2088) in CA1 neurons from mice treated with control IgG (n = 34 dendrites from 3 mice) and nivolumab (n = 42 dendrites from 3 mice). **(M)** Representative traces of outward A type potassium currents traces (left, middle) and quantification (amplitude, right) recorded from WT mice (WT: n = 17 neurons from 3 mice, WT + U0126: n = 20 neurons from 3 mice) and PD-1 KO mice (PD-1 KO : n = 18 neurons from 3 mice, PD-1 KO + U0126: n = 15 neurons from 3 mice). Note that the inhibitory effects were blocked by MEK inhibitor U0126. *P* = 0.0066 (WT vs. PD-1 KO); *P* = 0.0009 (PD-1 KO vs. PD-1 KO + U0126). Black trace: nonconditioned current; red trace: conditioned current. Bottom: protocol for isolation of A-type currents. A type current was isolated from the subtraction of red circle to black circle. **(N)** Representative traces of outward potassium A type current traces (left, middle) and amplitude (right) recorded from CA1 neurons treated with control IgG (IgG: n = 15 neurons from 3 mice, IgG + U0126: n = 12 neurons from 3 mice) and Nivolumab (Nivo: n = 14 neurons from 3 mice, Nivo + U0126: n = 11 neurons from 3 mice). Note that inhibitor effects were blocked by MEK inhibitor U0126. IgG vs. Nivolumab, *P* = 0.0075; Nivolumab vs. Nivolumab + U0126, *P* = 0.0100. Data are represented as mean ± SEM. Two-tailed Student’s t test (A, B, D, F, H, I, K, L), One-way ANOVA (M, N). Also see **Figure S3**.

Long-term potentiation (LTP) is an important form of synaptic plasticity underlying learning and memory (Martin et al., 2000) and can be induced by high-frequency stimulation (HFS) of Schaffer collaterals in the CA1 region of hippocampal slices (Shi et al., 2018). To test the involvement of PD-1 in this form of synaptic plasticity, we recorded LTP from brain slices of WT and PD-1 KO mice following HFS. This protocol, revealed by field excitatory postsynaptic potential (fEPSP) in CA1 region, showed that LTP was greatly enhanced in PD-1 KO slices compared to WT slices (Figures 3C and 3D); LTP was also greatly enhanced in WT slices incubated with nivolumab (300 ng/mL, 2 h) compared to control IgG (Figures 3E and 3F).

LTP is also associated with dendritic morphological changes in hippocampal CA1 pyramidal neurons (Matsuzaki et al., 2004). Loss of PD-1 resulted in increased dendritic spine density but unaltered spine size in CA1 neurons of PD-1 KO mice (Figures 3G-3I). Furthermore, anti-PD-1 treatment in WT slices with nivolumab (300 ng/mL, 2 h) increased spine density but not spine size (Figures 3J-3L). *Pdcd1* deficiency did not affect neuronal survival in the hippocampus (Figures S3A and S3B). Together, these results suggest that PD-1 deficiency in mice promotes basal synaptic transmission, LTP, and synaptogenesis in hippocampal neurons, contributing to the observed improvement in learning and memory.

PD-1 regulates the functions of immune cells and sensory neurons via the activation of tyrosine phosphatase SHP-1 (Chen et al., 2017; Hebeisen et al., 2013). Blockade of PD-1 in spinal cord dorsal horn neurons resulted in SHP-mediated hyper-phosphorylation of ERK (Jiang et al., 2020). To further explore the mechanism underlying the PD-1 regulation of synaptic plasticity, we examined the phosphorylation of ERK (P-ERK), as this process is critical for the induction of hippocampal LTP, enhancement of learning, and the hippocampus-dependent memory (Atkins et al., 1998; English and Sweatt, 1996, 1997; Sindreu et al., 2011). To test the hypothesis that PD-1 regulates learning and memory through ERK signaling, we examined P-ERK expression in hippocampal slices using immunohistochemistry following incubation of nivolumab or control IgG (300 ng/mL, 2 h). We observed marked induction of P-ERK in the dendrites of CA1 neurons after nivolumab incubation compared to control IgG (Figure S3C).

ERK was also shown to regulate neuronal excitability through the dendritic A-type K^+^ currents via direct phosphorylation of the Kv4.2 potassium channels (Adams et al., 2000; Schrader et al., 2006). Thus, we tested whether PD-1 could regulate neuronal excitability via altering A-type K^+^ currents. We used whole-cell, voltage-patch-clamp method to record the Kv4.2-mediated A-type currents in CA1 neurons of brain slices from WT and PD-1 KO mice. We observed a substantial reduction in the amplitude of A-type currents in PD-1 KO neurons compared to WT neurons; but this reduction was completely restored by the MEK (ERK kinase) inhibitor U0126 (Figure 3M). Furthermore, the reduced amplitude of A-type K^+^ currents was observed in nivolumab-treated WT CA1 neurons, in an ERK-dependent manner (Figure 3N). These results indicate that ERK-mediated suppression of A-type K^+^ currents in CA1 neurons promotes neuronal hyperactivity after loss or pharmacological inhibition of PD-1 (Figure S3D).

### PD-1 regulates neuronal excitability in non-human primate hippocampal neurons

To demonstrate the translational relevance of our findings, we tested the role of PD-1 in hippocampal neurons of non-human primates (NHP). *In situ* hybridization revealed expression of *PDCD1* mRNA in many hippocampal CA1 and CA3 neurons from Rhesus macaque monkey; quantitative analysis showed *PDCD1* mRNA expression by 70%-80% CA1 and CA3 neurons (Figures 4A and 4B). Notably, this percentage of positive neurons was much higher in NHP than that in mice (Figure S2D). The specificity of *PDCD1* staining was tested by the negative-control probe (Figure S3E).

**Figure 4.**
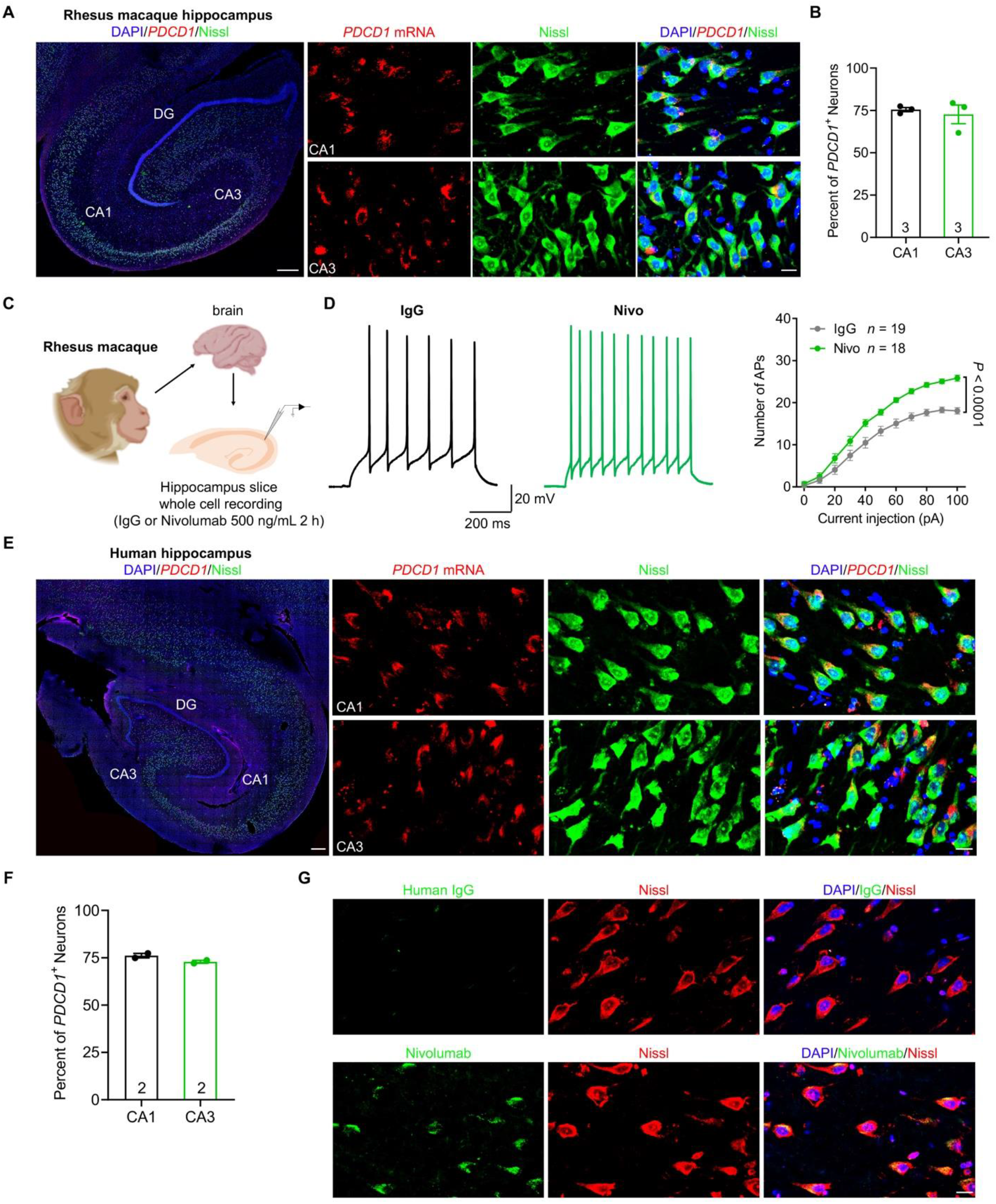
*PDCD1* is widely expressed by NHP and human hippocampal neurons. **(A)** *PDCD1* mRNA expression in Rhesus macaque hippocampal neurons as shown by *in situ* hybridization. Left, representative whole hippocampus images (Scale bar, 500 μm). Right panels, high magnification images showing *PDCD1* expression in Nissl-labeled neurons (Scale bar, 20 μm). **(B)** Quantification of *PDCD1*-postive neurons (75% of Nissl-labeled cells) in Rhesus macaque CA1 and CA3. n = 3 Rhesus macaques. **(C)** Schematic of experimental design for whole-cell patch clamp recordings in Rhesus macaque (NHP) hippocampal neurons. **(D)** Representative traces and quantification of current-evoked action potentials in CA1 neurons of brain slices treated with control IgG (n = 19 neurons from 3 NHPs) and nivolumab (n = 18 neurons from 3 NHPs). Nivolumab treatment increases neuronal excitability in Rhesus macaque hippocampal neurons (*P* < 0.0001). **(E)** *PDCD1* mRNA expression in human hippocampal neurons as shown by *in situ* hybridization. Left, representative whole hippocampus image (Scale bar, 500 μm). Right panels, high magnification images showing *PDCD1* expression in Nissl-labeled neurons (Scale bar, 20 μm). **(F)** Quantification of *PDCD1*-postive neurons in human CA1 and CA3. n = 2 human donors. **(G)** Representative images show CA1 neurons binding to nivolumab (human anti-PD-1 antibody) but not to control IgG. Scale bar, 20 μm. Data are represented as mean ± SEM. Two-way ANOVA (D). Also see **Figure S3**.

We also conducted patch-clamp recordings from 3 NHPs (Rhesus macaque, 2 females and 1 male at age of 13, 10, and 19 years). We prepared brain slices (350-μm thick) from NHP within 2 h of euthanasia. We treated the brain slices with control IgG or nivolumab treatment (500 ng/mL) for 2 h before recordings (Figure 4C). We induced action potentials by current injection (0-100 pA) and found that, compared with IgG control, nivolumab incubation significantly increased action potential-firing number evoked by current injection (*P* < 0.0001, Nivo vs. IgG control, n = 19 or 18 neurons, Figure 4D).

We also investigated the expression of *PDCD1* in the human hippocampal sections from 2 donors and found widespread expression of *PDCD1* mRNA in majority of CA1 and CA3 neurons (Figure 4E); RNAscope revealed that 76% of CA1 neurons and 72% of CA3 neurons express *PDCD1* (Figure 4F). The specific staining of *PDCD1* was also validated by a negative-control probe (Figure S3F). In addition, we performed PD-1 immunostaining in the human hippocampus and observed that nivolumab, but not human IgG, was able to bind CA1 neurons (Figure 4G). Collectively, these data reveal that PD-1 is extensively expressed by hippocampal neurons of different species, including mouse, monkey and human, and PD-1 down-regulates the function of mouse and NHP CA1 neurons.

### Conditional deletion of *Pdcd1* in excitatory neurons but not microglia improves learning and memory

To directly investigate the neuron-specific role of PD-1 in synaptic functions and memory, we generated *Pdcd1*-floxed mice (*Pdcd1^fl/fl^*) by flanking exon 2 with loxP sites (Figure 5A and Figure S4); this exon encodes a critical function (Keir et al., 2007). We used two strategies to test the function of neuronal PD-1 in neurons. First, we did bilateral stereotactic injections of an AAV-*Camk2a*-mCherry:Cre virus into the hippocampus of *Pdcd1^fl/fl^* mice (Figure S5A), which lead to robust AAV expression in CA1 and CA3 neurons (Figure S5B) and specific depletion of PD-1 expression in hippocampal excitatory neurons (Figure S5C). Mice that received AAV-*Camk2a*-mCherry:Cre injections exhibited enhanced learning and memory in the NOR test (Figure S5D). The hippocampus-specific knockout of PD-1 phenocopied the effects of global knockout in the MWM tests, resulting in improved spatial learning and memory in the probe tests but no change in swimming speed (Figures S5E-S5H).

**Figure 5.**
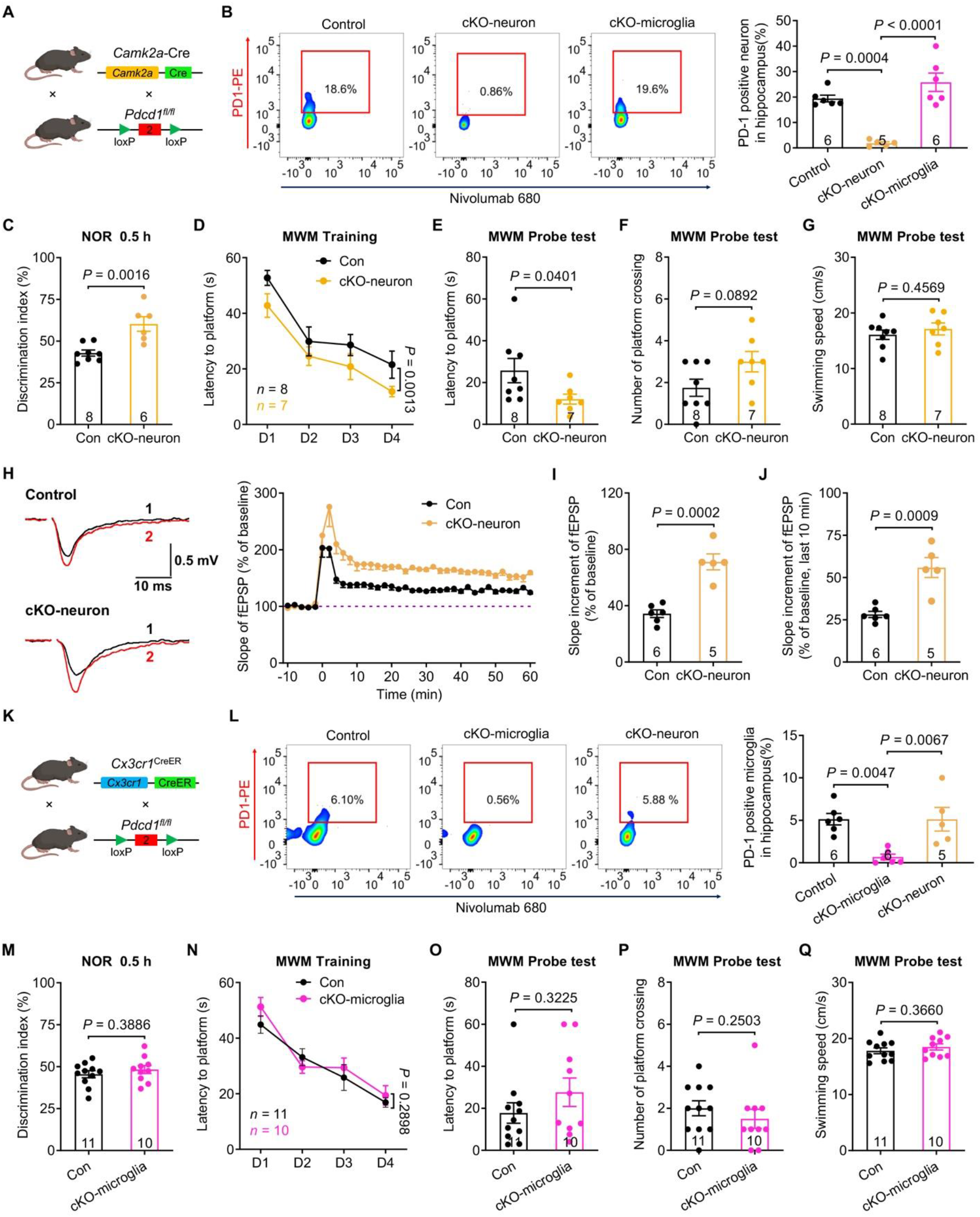
Conditional knockout of PD-1 in excitatory neurons but not microglia improves memory and synaptic plasticity. **(A)** Schematic illustration of the strategy to delete *Pdcd1* in excitatory neurons in conditional knockout (cKO-neuron) mice. **(B)** Flow cytometry shows the percentage of PD-1^+^ neurons in the hippocampi of cKO-neuron mice (n = 5), control littermates (n = 6), cKO-microglia mice, (n = 6). Note that PD-1 is lost in CaMKIIα^+^ neurons in cKO-neuron mice but not in cKO-microglia mice. *P* = 0.0004 vs. control littermates; *P* < 0.0001 vs. cKO-microglia mice. **(C)** NOR test in control littermates (n = 8) and cKO-neuron mice (n = 6). Note that cKO-neuron mice exhibit higher discrimination index (*P* = 0.0016). **(D)** Spatial learning curve during MWM training in control littermates (n = 8) and cKO-neuron mice (n = 7). cKO-neuron mice spend less time locating the hidden platform compared to control littermates (*P* = 0.0013). **(E-F)** MWM probe test for latency to platform (E) and number of platform crossings (F) in control littermates (n = 8) and cKO-neuron mice (n = 7). cKO-neuron mice have shorter latency (*P* = 0.0401) and slightly increased platform zone crossings number (*P* = 0.0892). **(G)** In MWM test control littermates (n = 8) and cKO-neuron mice (n = 7) are comparable in swimming speed (*P* = 0.4569). **(H)** Representative traces (left) and summary plots (right) of LTP induced by 2 × HFS in CA1 region of control littermates and cKO-neuron slices Left: Black traces (1) and red trace (2) represent the baseline fEPSP and post-induction fEPSP, respectively. **(I-J)** Quantification of average slope of LTP in CA1 region of control littermate slices (n = 6 from 5 mice) and cKO-neuron slices (n = 5 from 5 mice) both in entire phase (I, *P* = 0.0002) and late phase (J, *P* = 0.0009). **(K)** Schematic illustration of the strategy to delete *Pdcd1* in microglia in conditional knockout (cKO-microglia) mice. **(L)** Flow cytometry shows the percentage of PD-1^+^ microglia in the hippocampi of cKO-microglia mice (n = 6), control littermates (n = 6), cKO-neuron mice (n = 5). Note that PD-1 is lost in Cx3cr1^+^ microglia in cKO-microglia mice but not in cKO-neuron mice. *P* = 0.0047 vs. control littermates; *P* = 0.0067 vs. cKO-neuron mice. **(M)** NOR test shows comparable discrimination index in control littermates (n = 11) and cKO-microglia mice (n = 10, *P* = 0.3886). **(N)** Spatial learning curves during MWM training show comparable learning capacity between control littermates (n = 11) and cKO-microglia mice (n = 10, *P* = 0.2898). **(O-P)** MWM probe test for latency to platform (O) and platform zone crossings number (P), showing no difference (O, *P* = 0.3225; P, *P* = 0.2503) between control littermates (n = 11) and cKO-microglia mice (n = 10). **(Q)** MWM probe test shows comparable swimming speed (*P* = 0.3660) in control littermates (n = 11) and cKO-microglia mice (n = 10). Data are represented as mean ± SEM. One-way ANOVA (B, L), Two-tailed Student’s t test (C, G, I, J, M, Q), Mann-Whitney test (E, F, O, P), Two-way ANOVA (D, N). Also see **Figures S4-S6**.

Next, we crossed the *Pdcd1^fl/fl^* mice with *Camk2a-Cre* mice to generate conditional knockout of *Pdcd1* in excitatory neurons, described as cKO-neuron (Figure 5A). For comparison, we crossed the *Pdcd1^fl/fl^* mice with *Cx3cr1^CreER^* mice to generate conditional knockout of *Pdcd1* in microglia and monocytes, described as cKO-microglia. Flow cytometry analysis revealed that the percentage of PD-1^+^ excitatory neurons was substantially decreased in the hippocampus of cKO-neuron mice, suggesting a loss of PD-1 in hippocampal excitatory neurons (Figure 5B and Figure S6). Genotyping and *in situ* hybridization validated a loss of *Pdcd1* mRNA expression in CA1 neurons of the cKO-neuron mice (Figures S5I and S5J). cKO-neuron mice displayed greater object learning and memory capacity than control littermates (Figure 5C). They also showed substantially enhanced learning in the MWM hidden platform training and improved memory in the probe tests but unaltered swimming speed (Figures 5D-5G). The cKO-neuron mice had normal motor function and general emotional states (Figures S5K-S5O). We recorded LTP in hippocampal neurons from brain slices of cKO-neuron and littermate control mice following HFS. The cKO mice had improved LTP compared to controls; as indicated by summary plots and average slope of fEPSP during the entire 60 min (Figure 5I) and just the last 10 min (Figure 5J). These results confirm the observations with the global KO mice and reveal that PD-1 effects on learning and memory are largely restricted to the hippocampus and excitatory neurons.

Studies have shown that the PD-1 is expressed by microglia and may play a role in neurodegenerative diseases (Suda et al., 2021; Tabula Muris et al., 2018; Xing et al., 2021; Yao et al., 2014; Zeisel et al., 2018). However, the physiological role of PD-1 in microglia/monocytes in cognition remains unclear. FACS analysis in cKO-microglia mice showed that the percentage of PD-1^+^ microglia was greatly decreased in the hippocampus as compared with that in control littermate mice or the cKO-neuron mice (Figures 5K and 5L and Figure S6). Genotyping and *in situ* hybridization validated the loss of *Pdcd1* mRNA in hippocampus of cKO-microglia mice (Figures S5P and S5Q). Furthermore, cKO-microglia mice and control littermates displayed comparable learning and memory performance in NOR and MWM tests (Figures 5M-5P), as well as comparable swimming speed (Figure 5Q), motor function, and general emotional states (Figures S5R-S5V). Together, these data suggest that PD-1 in microglia/monocytes does not impact cognition under these testing conditions.

### Selective PD-1 re-expression in hippocampal excitatory neurons of PD-1 KO mice impairs memory and synaptic plasticity

Our findings showed that loss of PD-1 in hippocampal excitatory neurons, but not microglia, plays a crucial role in memory enhancement. To investigate whether re-expression of PD-1 in the hippocampus of PD-1 KO mice would be sufficient to reverse cognitive changes, we constructed AAV expressing either *Pdcd1* driven by *Camk2a* promoter (AAV-*Pdcd1*) or control fluorescent protein eGFP (AAV-control) and delivered the AAV specifically to the hippocampus by bilateral stereotactic injections (Figure 6A). This strategy resulted in selective re-expression of PD-1 in the targeted region including CA1 and CA3 neurons (Figure 6B and Figures S7A and S7B). *In situ* hybridization analysis confirmed the expression of *Pdcd1* in hippocampal CA1 and CA3 neurons (Figures 6C and 6D). No neuronal loss was observed in the hippocampus after the re-expression (Figures S7C). Four weeks after the AAV infection, we conducted cognitive behavioral tests and recorded LTP in hippocampal slices. Viral expression of PD-1 in the KO mice retarded object learning and memory capacity in NOR test (Figure 6E) and impaired the learning and memory performance in MWM tests (Figures 6F-6I), with no effects on motor activities or anxiety and depression-like behaviors (Figures S7D-S7H). PD-1 re-expression in KO background also suppressed LTP (Figure 6J). Notably, the average slope of fEPSP in the viral-infected slices were markedly reduced, especially in the late-phase (Figures 6K and 6L). Thus, hippocampal neuronal rescue of PD-1 expression is sufficient to reverse both the cognitive and synaptic changes in PD-1 KO mice, supporting the critical role of neuronal PD-1 for the control of memory and synaptic plasticity.

**Figure 6.**
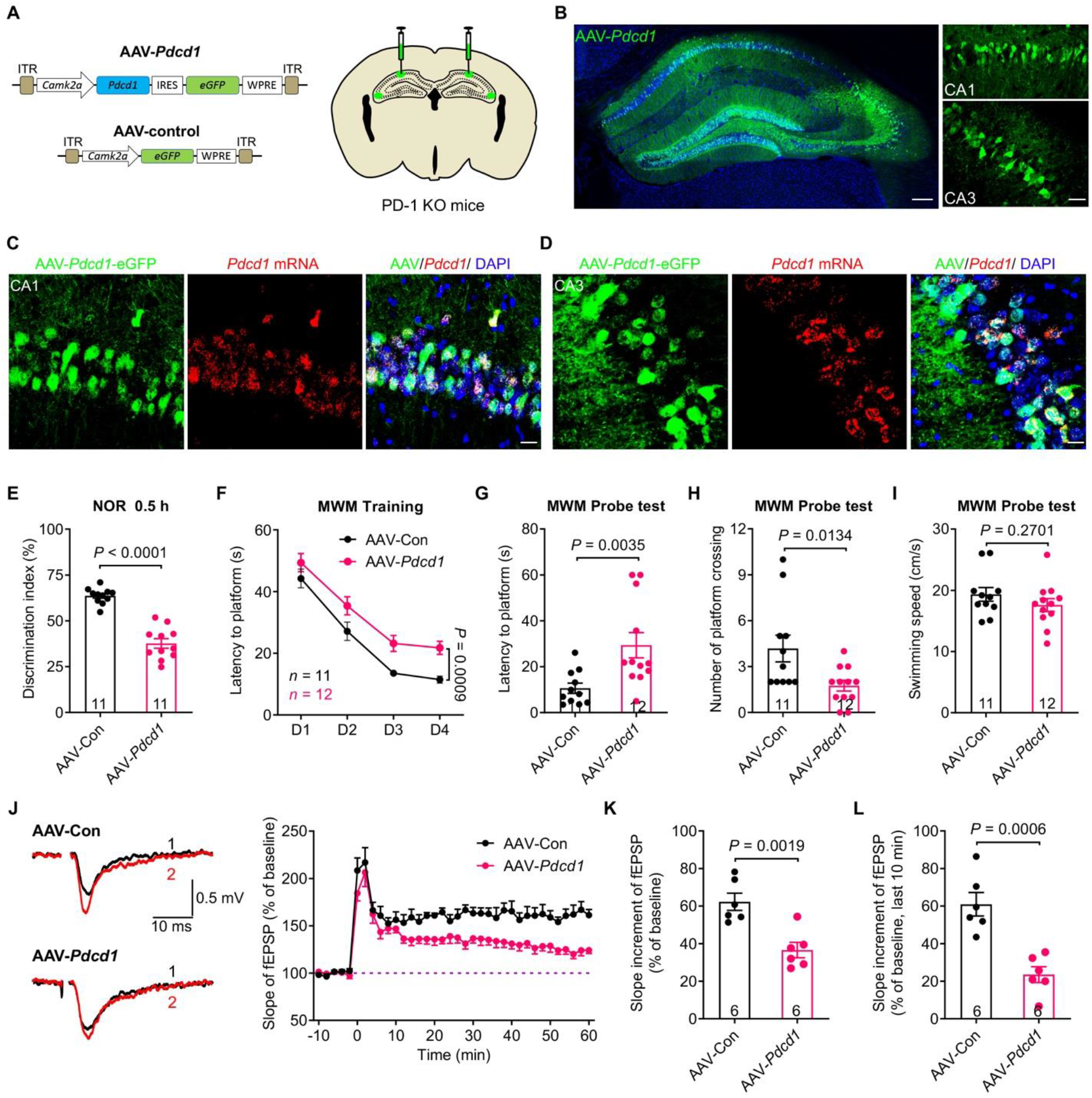
Selective PD-1 re-expression in the hippocampus of PD-1 KO mice impairs memory and synaptic plasticity. **(A)** Schematics of AAV constructs of *Pdcd1* (AAV-Pdcd1) or control (AAV-control) (left) and bilateral viral injections into the mouse hippocampus (right). Abbreviations: *Camk2a*, Ca^2+^/calmodulin-dependent protein kinase II alpha prompter; ITR, inverted terminal repeats; IRES, internal ribosome entry site; WPRE, woodchuck hepatitis virus posttranscriptional regulatory element. **(B)** Representative confocal images of mouse hippocampus (left) and infected CA1 and CA3 neurons (right) after the AAV injections. Blue: DAPI. Scale bars, 200 µm (left), 50 µm (right). **(C-D)** Representative confocal images of the infected CA1 (C) and CA3 (D) neurons co-expressing eGFP (AAV-*Pdcd1*) and *Pdcd1* mRNA (revealed by RNAscope). Scale bar, 20 µm. **(E)** NOR test in PD-1 KO mice injected with AAV-control (n = 11) and AAV-*Pdcd1* (n = 11). Note that mice injected with AAV-*Pdcd1* show deficits in NOR test (*P* < 0.0001). **(F)** Spatial learning curves during MWM training in PD-1 KO mice injected with AAV-control (n = 11) and AAV-*Pdcd1* (n = 12), measured as latency to find the hidden platform. PD-1 KO mice injected with AAV-*Pdcd1* show deficits (*P* = 0.0009). **(G-H)** MWM probe test shows deficits of PD-1 KO mice injected with AAV-*Pdcd1* (n = 12) vs. AAV-control (n = 11) in locating the platform (G, *P* = 0.0035) and crossing the platform zone (H, *P* = 0.0134). **(I)** MWM probe test shows comparable swimming speed (*P* = 0.2701) in PD-1 KO mice injected with AAV-control (n = 11) and AAV-*Pdcd1* (n = 12). **(J)** Representative traces (left) and summary plots (right) of LTP induced by 2 × HFS in the CA1 region of PD-1 KO mice injected with AAV-control or AAV-*Pdcd1*. Left: Black traces (1) and red trace (2) represent the baseline fEPSP and post-induction fEPSP, respectively. **(K-L)** Quantification of average slope of LTP in the CA1 region of PD-1 KO mice injected with AAV-control (n = 6 slices from 5 mice) and AAV-*Pdcd1* (n = 6 slices from 6 mice). Note that PD-1 KO mice injected with AAV-*Pdcd1* show deficits in generating LTP both in entire phase (K, *P* = 0.0019) and late phase (L, *P* = 0.0006). Data are represented as mean ± SEM. Two-tailed Student’s t test (E, I, K, L), Mann-Whitney test (G, H), Two-way ANOVA (F). Also see **Figure S7**.

### Suppression of PD-1 improves cognitive decline after traumatic brain injury

Traumatic brain injury (TBI) causes significant cognitive decline (Harrison et al., 2015; Wang et al., 2007). To define the role of PD-1 in this condition, we used the closed-head TBI model in mice (Wang et al., 2007) (Figure 7A). We established the TBI model in WT and PD-1 KO mice (10-12 weeks old) and tested their motor coordination to determine the time course of motor recovery after TBI. Rotarod testing revealed similar motor impairment in both WT and PD-1 KO mice, which improved during the next 3 weeks (Figure 7A). We conducted the NOR test 3 weeks after TBI in WT and PD-1 KO that had full motor recovery (Figure 7B). Compared with sham surgery, TBI caused a lower discrimination index in WT mice, but PD-1 KO mice had a markedly higher discrimination index than control mice, indicating *Pdcd1* deletion protects against the TBI-induced cognitive decline (Figure 7B). In MWM hidden platform training, TBI resulted in a cognitive deficit in WT mice, but PD-1 KO mice were protected from TBI, showing an identical learning curve to WT mice without TBI (sham surgery, Figure 7C). In the MWM probe tests, PD-1 KO mice located the platform zone much more quickly (Figure 7D), crossed the platform zone frequently and had a higher swimming speed compared to control WT mice with TBI (Figures 7E and 7F). These results indicate that deletion of PD-1 confers protection against the cognitive defects after TBI.

**Figure 7.**
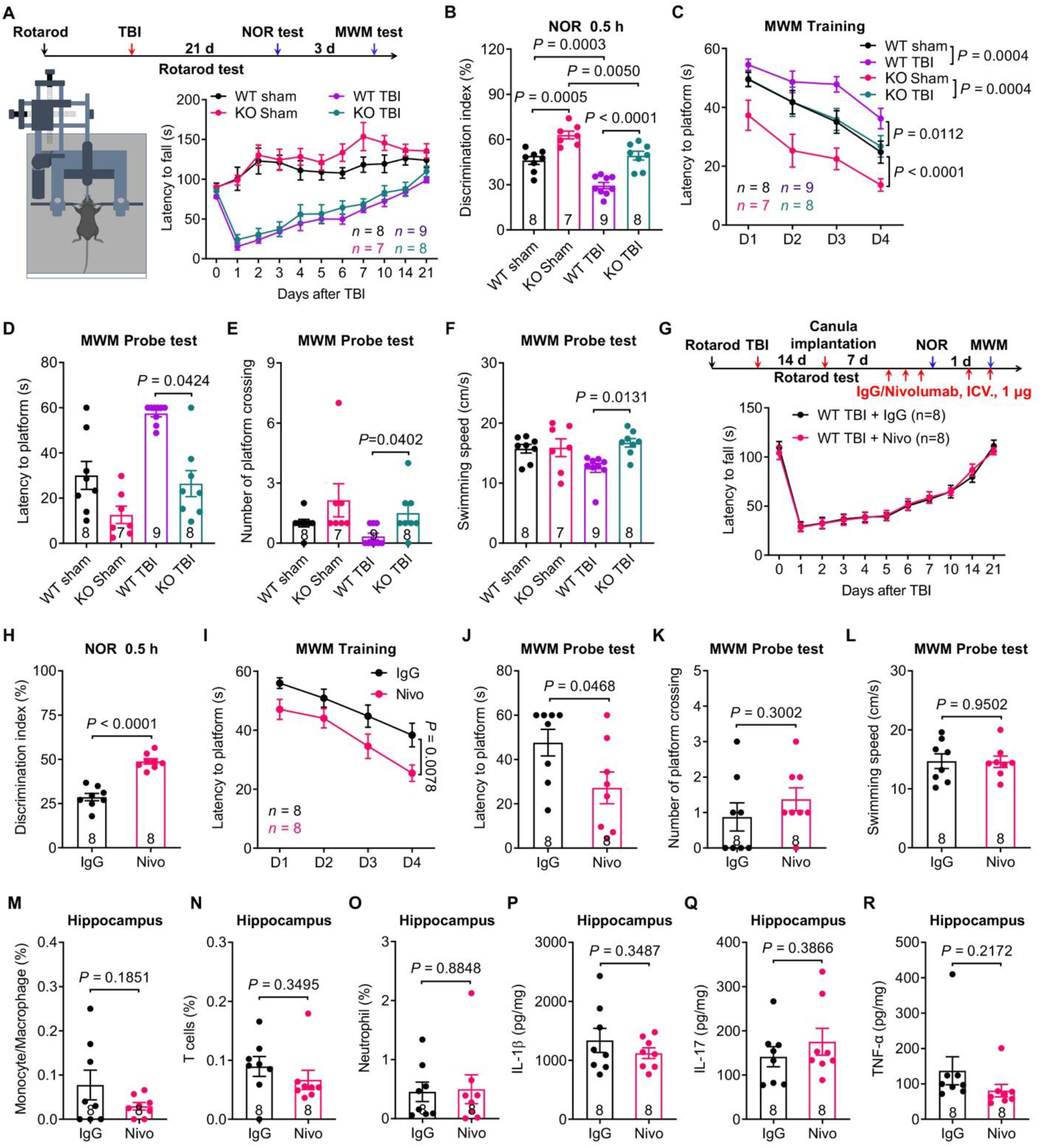
*Pdcd1* deficiency or ICV blockade of PD-1 protects cognitive function after traumatic brain injury. **(A-F)** Rotarod test (A), NOR test (B), and MWM training test (C) and probe tests (D-F) in WT and PD-1 KO mice with traumatic brain injury (TBI, WT: n = 9, PD-1 KO : n = 8) or sham surgery (WT: n = 8, PD-1 KO : n = 7). **(A)** Experimental paradigm and rotarod test in WT mice and PD-1 KO mice with TBI or sham surgery. WT and PD-1 KO mice show comparable motor impairment that recovers after 3 weeks. **(B)** NOR testing shows cognitive deficits in both WT TBI mice (*P* = 0.0003, vs. Sham) and PD-1 KO TBI mice (*P* = 0.0050, vs. Sham); but PD-1 KO TBI mice have a higher discrimination index than WT TBI mice (*P* < 0.0001). **(C)** MWM training curves show deficits in both WT (*P* = 0.0004 vs. Sham) mice and PD-1 KO (*P* = 0.0004, vs. Sham); but PD-1 KO TBI mice spend less time on navigating the hidden platform location than WT TBI mice (*P* = 0.0112). **(D-E)** MWM probe tests for latency to platform (D) and number of platform zone crossings (E). Note that PD-1 KO TBI mice exhibit shorter latency of locating the platform zone (D, *P* = 0.0424, vs. WT) and more platform zone crossings number (E, *P* = 0.0402, vs. WT). **(F)** MWM probe test shows higher swimming velocity in PD-1 KO TBI mice than WT TBI mice (*P* = 0.0131). **(G-R)** Rotarod test (G), NOR test (H), and MWM training test (I), probe test (J-L), and FACS (M-O) and ELISA (P-R) analyses in WT TBI mice treated with ICV injections of control IgG or nivolumab (1 μg per mouse, every other day). n = 8 mice per group. **(G)** Experimental paradigm for rotarod test (day 1-21), drug injections (day 22, 24, 26, 28, 30, 32, 34), and NOR test (day 27, 28), MWM test (day 30, 31, 32, 33, 34), and tissue collection (day 35). **(H)** NOR test in TBI mice treated with control IgG and nivolumab. Note that nivolumab treated TBI mice have significant improvement in object learning and memory (*P* < 0.0001). **(I)** Spatial learning curves during MWM training show significant improvement (*P* = 0.0078) in WT TBI mice treated with nivolumab than control IgG. **(J-K)** MWM probe test for latency to platform (J) and number of platform crossings (K) in control IgG and nivolumab treated mice. Nivolumab treated mice exhibit a shorter latency to locate the platform zone (*P* = 0.0468) but comparable platform zone crossings number (*P* = 0.3002). **(L)** MWM probe test shows comparable swimming speed (*P* = 0.9502) between control IgG and nivolumab treated TBI mice. **(M-O)** Flow cytometry analysis of hippocampus tissues from TBI mice treated with control IgG and nivolumab analyzed for the percentage of immune cells (M: monocyte/macrophage, *P* = 0.1851; N: T cells *P* = 0.3495; O: neutrophils, *P* = 0.8848). **(P-R)** ELISA test of hippocampus tissues from TBI mice treated with control IgG and nivolumab analyzed for the level of cytokines (P: IL-1β, *P* = 0.3487; Q: IL-17, *P* = 0.3866; R: TNF-α, *P* = 0.2172). Data are represented as mean ± SEM. One-way ANOVA (B, F), Kruskal-Wallis test (D, E), Two-way ANOVA (C, I), Two-tailed Student’s t test (H, L, M, N, O, P, Q, R), Mann-Whitney test (J, K). Also see **Figure S8**.

To further demonstrate whether anti-PD-1 treatment could improve cognitive deficits after TBI, we administered nivolumab or control IgG via ICV injections to block the PD-1 function in the brain. Cognitive testing was conducted after animal recovery from motor deficits and three injections of nivolumab or control IgG (3 × 1 µg, every other day, Figure 7G). Nivolumab-treated TBI mice exhibited marked improvements in learning and memory capacity in the NOR test (Figure 7H) and spatial learning in the MWM training and probe tests (Figures 7I and 7J) with unaltered platform crossing number and swimming activity (Figures 7K and 7L). Since anti-PD-1 treatment is a well-utilized immunotherapy strategy, we examined changes in immune cell types and pro-inflammatory cytokines in hippocampus and cortex tissues after nivolumab or IgG control treatment in TBI mice. Flow cytometry showed comparable abundance of immune cell types, including monocytes/macrophages, T cells and neutrophils after nivolumab or control IgG treatment in the hippocampus (Figures 7M-7O) and cortex (Figures S8A-S8E). The levels of pro-inflammatory cytokines in the hippocampus and cortex, including IL-1β, IL-17 and TNF-α, were also comparable between control IgG and nivolumab treatment (Figures 7P-7R and Figures S8F-S8H). These results suggest that CNS anti-PD-1 treatment improved cognitive deficits after TBI, through a mechanism that does not appear to recruit immune cells to the brain (Figure S3D).

## DISCUSSION

We have revealed a previously unrecognized physiological role of PD-1 as a neuromodulator that regulates neuronal activity and excitability, leading to altered cognitive function. We provided several lines of evidence supporting this observation. First, loss of PD-1 in global knockout mice enhanced learning and memory in both NOR and MWM tests and increased LTP in hippocampal CA1 neurons. Second, using a conditional knockout mice with selective *Pdcd1* deletion in excitatory neurons (cKO-neuron), we observed similar behavioral and cellular phenotypes as in the global knockout mice. Consistently, injections of AAV-*Camk2a*-mCherry:Cre virus in the hippocampus of *Pdcd1^fl/fl^* mice enhanced cognition, whereas re-expression of *Pdcd1* in hippocampal neurons via AAV-*Pdcd1* injection in the global knockout mice suppressed cognition. In contrast, cKO mice lacking *Pdcd1* in microglia and monocytes (cKO-microglia) showed no changes in the cognitive tests. Third, intracerebroventricular administration of nivolumab, a clinically used anti-PD-1 immunotherapy, enhanced learning and memory in adult WT mice and further improved cognitive function in mice with TBI. Perfusion of brain slices with nivolumab was sufficient to enhance LTP, possibly through ERK-mediated suppression of A-type potassium currents in CA1 neurons. Fourth, neuronal-PD1 signaling is conservative across different species, as *PDCD1* mRNA and PD-1 are extensively expressed in NHP and human CA1 neurons, and nivolumab was sufficient to increase action potential firing in CA1 neurons of NHP.

PD-1 is a well-known immune checkpoint inhibitor, which is the foundation of the anti-PD-1 based immunotherapies that have achieved success in treating some cancer patients (Butte et al., 2007; Freeman et al., 2000; Keir et al., 2008; Nishimura et al., 1999; Sharma and Allison, 2015). Recent studies have shown that PD-1 is widely expressed in various cell types, including T cells, macrophages, microglia, and melanoma cells (Gordon et al., 2017; Kleffel et al., 2015; Wang et al., 2020a; Yao et al., 2014). Increasing evidence suggests PD-1 is expressed by neurons, including retinal ganglion neurons (Sham et al., 2012), primary sensory neurons of dorsal root ganglion (Chen et al., 2017; Liu et al., 2020), and spinal cord, thalamic, and cortical neurons (Jiang et al., 2020). Single-cell RNA-seq revealed low-level of *Pdcd1* mRNA in cortical and hippocampal neurons (Tabula Muris et al., 2018; Zeisel et al., 2018). Single-cell RNA-seq may detect ony the most abundant mRNAs (Islam et al., 2014). Using more sensitive RNAscope *in situ* hybridization, our results showed that *Pdcd1* mRNA is present in 55-65% mouse CA1/CA3 neurons. Notably, the percentage of *PDCD1* mRNA^+^ neurons is higher in NHP and human CA1/CA3 neurons (70-80%), suggesting a more active role of neuronal PD-1 in higher species. Since cellular concentrations of proteins and the abundance of their corresponding mRNAs are not necessarily correlated (Vogel and Marcotte, 2012), it was essential for us to observe specific PD-1 protein expression in hippocampal CA1 and CA3 neurons of mouse, NHP, and human tissues. Most importantly, we demonstrated the functional neuronal PD-1 at both cellular and behavioral levels.

At the cellular levels, loss of PD-1 resulted in increased neuronal excitability in CA1 neurons, and strikingly, 40% of *Pdcd1-*deficient CA1 neurons exhibited spontaneous discharges. PD-1-mediated excitability change could dirve the synaptic changes observed in this study. Loss of PD-1 resulted in enhanced synaptic plasticity, as indicated by increased excitatory synaptic transmission (EPSC) and increased slope of LTP, associated with increased number of dendritic spines. These changes appear to be neuron intrinsic, as dissociated neurons in primary cultures of PD-1 KO mice also showed increased excitability. However, neuronal survival was not altered in PD-1 KO mice. Intriguingly, inhibition of PD-1 with nivolumab at very low concentration (7 nM) in WT neurons is able to produce similar cellular phenotypes of PD-1 KO mice, including increases in neuronal excitability, LTP slope, and number of dendritic spines. Perfusion of nivolumab but not control IgG also increased excitability of CA1 neurons of NHP, supporting an active role of neuronal PD-1 signaling in primates.

Mechanistically, our results suggest that PD-1-mediated ERK phosphorylation and A-type K^+^ current modulation may underlie neuronal and synaptic changes in CA1 neurons after PD-1 deficiency (Figure S3D). Tyrosine phosphatase SHP-1 was implicated in mediating the PD-1’s biological actions in immune cells (Hebeisen et al., 2013) and primary sensory neurons (Chen et al., 2017; Liu et al., 2020). Blockade of PD-1 in spinal cord slices resulted in SHP-mediated hyper-phosphorylation of ERK in spinal cord neurons (Jiang et al., 2020). ERK activation by phosphorylation is essential for the induction of hippocampal LTP and the formation of the hippocampus-dependent memory (Atkins et al., 1998; English and Sweatt, 1996, 1997; Sindreu et al., 2011). Our data showed that PD-1 blockade with nivolumab is sufficient to cause downstream signaling by increasing ERK phosphorylation in CA1 neurons. ERK enhances learning and memory by inducing dendritic spine formation and stabilization as well as CREB-mediated transcription (Impey et al., 1999; Impey et al., 1998; Sweatt, 2004; Tang and Yasuda, 2017). ERK also regulates neuronal excitability through dendritic A-type K^+^ currents via direct phosphorylation of the Kv4.2 potassium channels in neurons (Hu et al., 2006; Schrader et al., 2006). Deletion of the gene encoding Kv4.2 eliminates dendritic A-type K^+^ current and enhances LTP induction in hippocampal CA1 neurons (Chen et al., 2006). A-type K^+^ currents are at high density in CA1 dendrites where the neurons receive synaptic input, and rapid voltage-dependent activation of these channels can limit the peak amplitude of back-propagating action potentials and exert profound effects on hippocampal synaptic functions and memory (Adams et al., 2000; Chen et al., 2006; Hoffman et al., 1997). Importantly, we observed increased P-ERK expression selectively in the dendrites of CA1 neurons after nivolumab incubation. In CA1 neurons we also found marked reductions in Kv4.2-mediated A-type currents after PD-1 deficiency or nivolumab-treatment, which can be reversed by ERK inhibition. Thus, we postulate that anti-PD-1 treatment-mediated activation ERK and inhibition of A-type K^+^ currents in CA1 neurons can cause the cellular and behavioral phenotypes that we observed in PD-1 KO mice and nivolumab-treated brain slices and animals. Additionally, anti-PD1-mediated inhibition of GABAergic signaling may also be responsible for neuronal hyperactivity (Jiang et al., 2020).

At behavioral levels, we employed loss-of-function approaches in neuron-specific cKO mice and AAV-treated mice with selective neuronal infection to demonstrate that lack of neuronal PD-1 is sufficient to enhance cognitive function. Conversely, our gain-of-function approach through PD-1 re-expression in PD-1-deficient hippocampal neurons impaired memory. By contrast, cognitive behaviors were unaltered in cKO mice lacking PD-1 in microglia and monocytes. Our data strongly suggest a physiological role of PD-1 in cognition through direct neuronal modulation. Furthermore, we found a marked improvement of cognitive function by intraventricular treatment of nivolumab in a mouse model of TBI. We did not see significant changes in immune cell types or pro-inflammatory cytokines in the hippocampus and cortex of the TBI-mice treated with nivolumab, suggesting that cognitive improvement results from neuronal modulation rather than recruitment of the immune cells (Figure S3D). Previous studies showed that systemic anti-PD-1 treatment enhanced memory in mouse AD models, via increasing the CNS recruitment of immune cells (e.g., macrophages) that can promote the clearance of amyloid-β plaques and alleviate tauopathy (Baruch et al., 2016; Rosenzweig et al., 2019; Xing et al., 2021). However, contradictory results were also reported showing no effective removal of amyloid-β plaques by anti-PD-1 treatments (Latta-Mahieu et al., 2018; Lin et al., 2019; Obst et al., 2018). While immunotherapy is appealing for treating AD, future studies are needed to test whether monotherapy is sufficient or polytherapy is required to increase “immunity” as in some cancer conditions (Sharma and Allison, 2015; Xing et al., 2021). Increasing evidence suggests that neuroinflammation after TBI and surgery contributes to cognitive decline, in part through production of pro-inflammatory cytokines (Salvador et al., 2021; Simon et al., 2017; Yang et al., 2020). Millions of cancer survivors suffer from neurological deficits including cognitive decline due to chemotherapy-induced microglial activation, neuroinflammation, and demyelination (Gibson and Monje, 2021; Gibson et al., 2019). Thus, immunotherapy could be both beneficial and detrimental for cognitive function in a context-dependent manner. Our results showed that intraventricular treatment with nivolumab for a short period (a few days) not only enhanced learning and memory in naïve animals but also restored cognition in mice with TBI. In nivolumab-treated TBI mice, we did not see changes in signs of neuroinflammation, including numbers of T cells and macrophages and expression of the pro-inflammatory cytokines TNF-α, IL-1b, and IL-17 in the cortex and hippocampus. However, we cannot rule out that systemic Nivolumab treatment may produce different effects on immune cell types and cytokines.

In summary, our findings support a “neurotherapy” of anti-PD-1 treatment for cognitive improvement, as PD-1 serves as a neuronal inhibitor and its inhibition can rapidly boost neuronal activity and enhance memory. However, chronic effects of anti-PD-1 treatment will certainly involve immune modulation, which can either impair memory via producing neuroinflammation or enhance memory via clearance of amyloid-β plaques under some neurodegenerative condition. Notably, nivolumab may reach to significant concentrations (35 to 150 ng/ml) in the cerebrospinal fluid in melanoma patients with metastasis to the brain (van Bussel et al., 2019). These concentrations may affect neuronal activities in the brain. It will be of great interest to study acute neuronal effects of anti-PD-1 therapies in cancer patients (e.g., patients with brain tumor) where significant CNS penetration of antibody could occur. Last but not least, expression of PD-1 in hippocampal excitatory neurons suppresses their activity under normal conditions, which dampens the cognitive ability of mice. This brake provides an opportunity for enhanced learning ability under special conditions by downregulating PD-1 function, which might occur by regulation of PD-1 levels or function (e.g., covalent modifications) in response to environmental signals. Activity of PD-1 might also be regulated by availability of its ligands (PD-L1 and PD-L2) or intracellular signaling molecules that it engages. These are prime areas for future research.

## ACKNOWLEDGMENTS

We thank Ji lab members and Katherine Peters for helpful discussion and Josh Huang for critical reading of the manuscript. This study was supported by Duke University Anesthesiology Research Funds. The NHP study was supported by NIH grant AR069861 to M.C.K.

## AUTHOR CONTRIBUTIONS

J.Z. and R.R.J developed the project. J.Z., S.B., A.M., K.F., C.J., A.R., C.D.R., Q.H., and H.W. conducted experiments and data analysis; R.D.P. generated *Pdcd1* floxed mice. M.C.K provided monkey hippocampus tissues. R.R.J. and J.Z. wrote the manuscript; R.D.P. and other co-authors edited the manuscript.

## DECLARATION OF INTERESTS

Dr. Ji is a consultant of Boston Scientific and received a research grant from the company for a project unrelated to this study. Other authors declare no completing interests.

## STAR★METHODS

### KEY RESOURCES TABLE

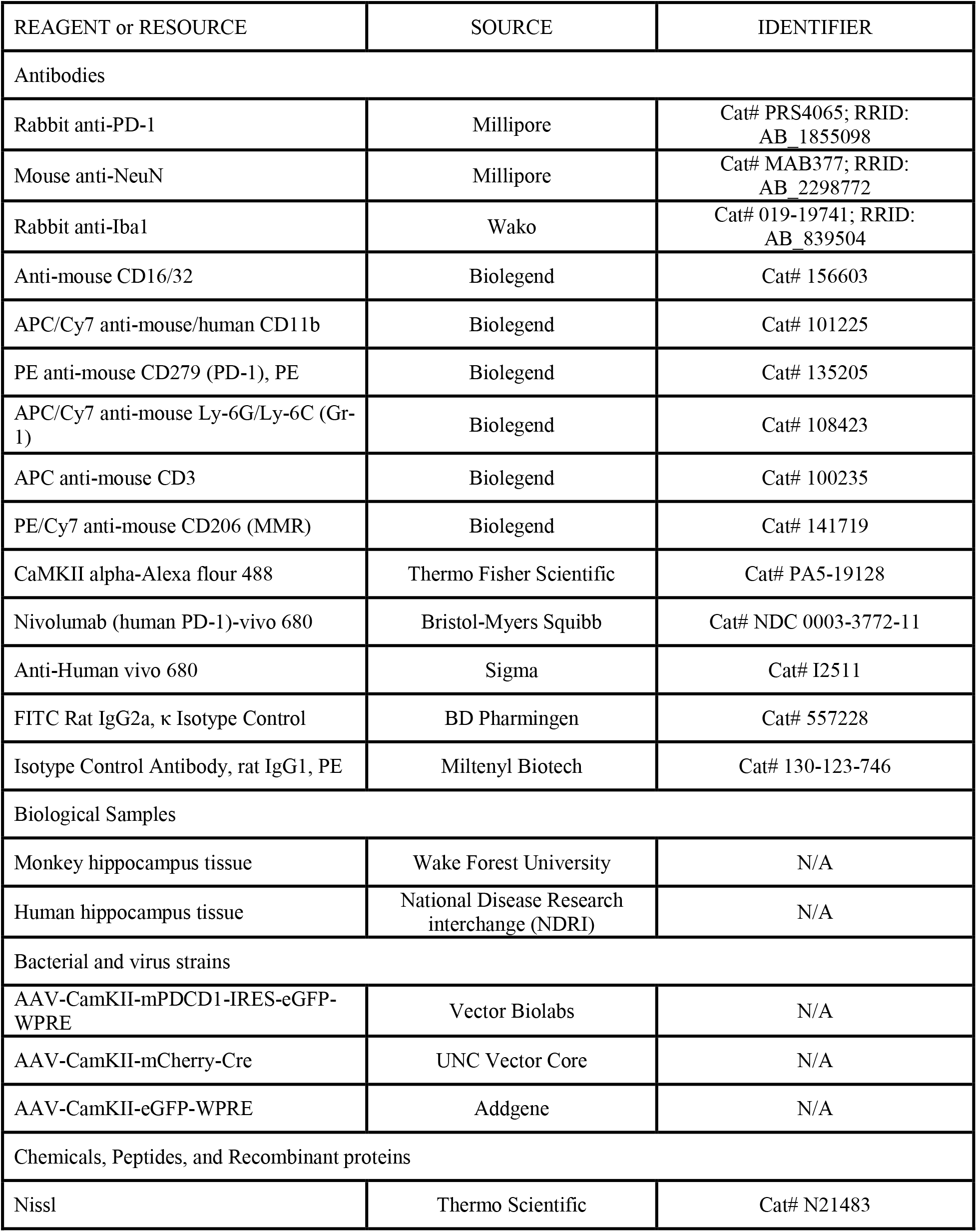
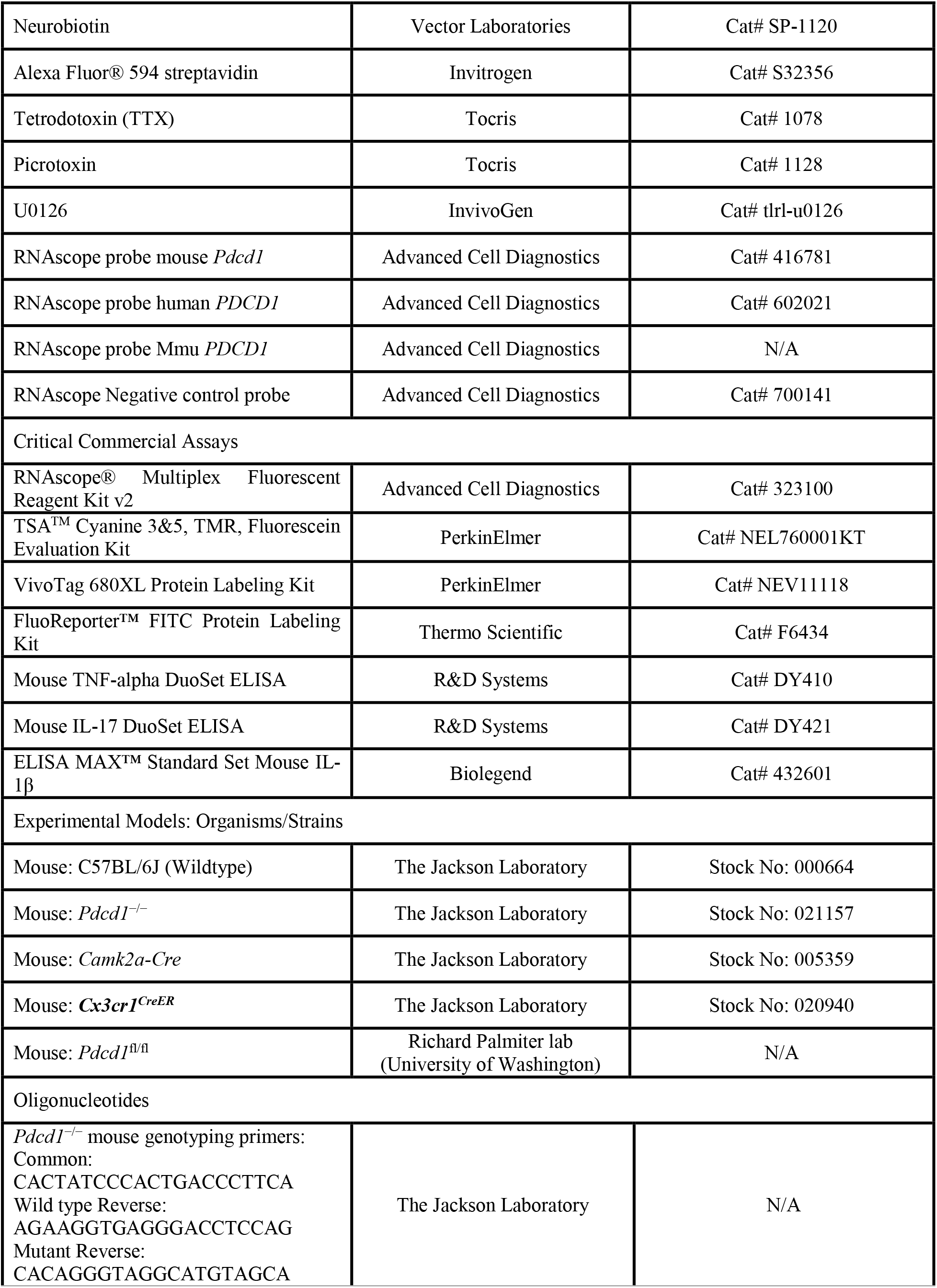
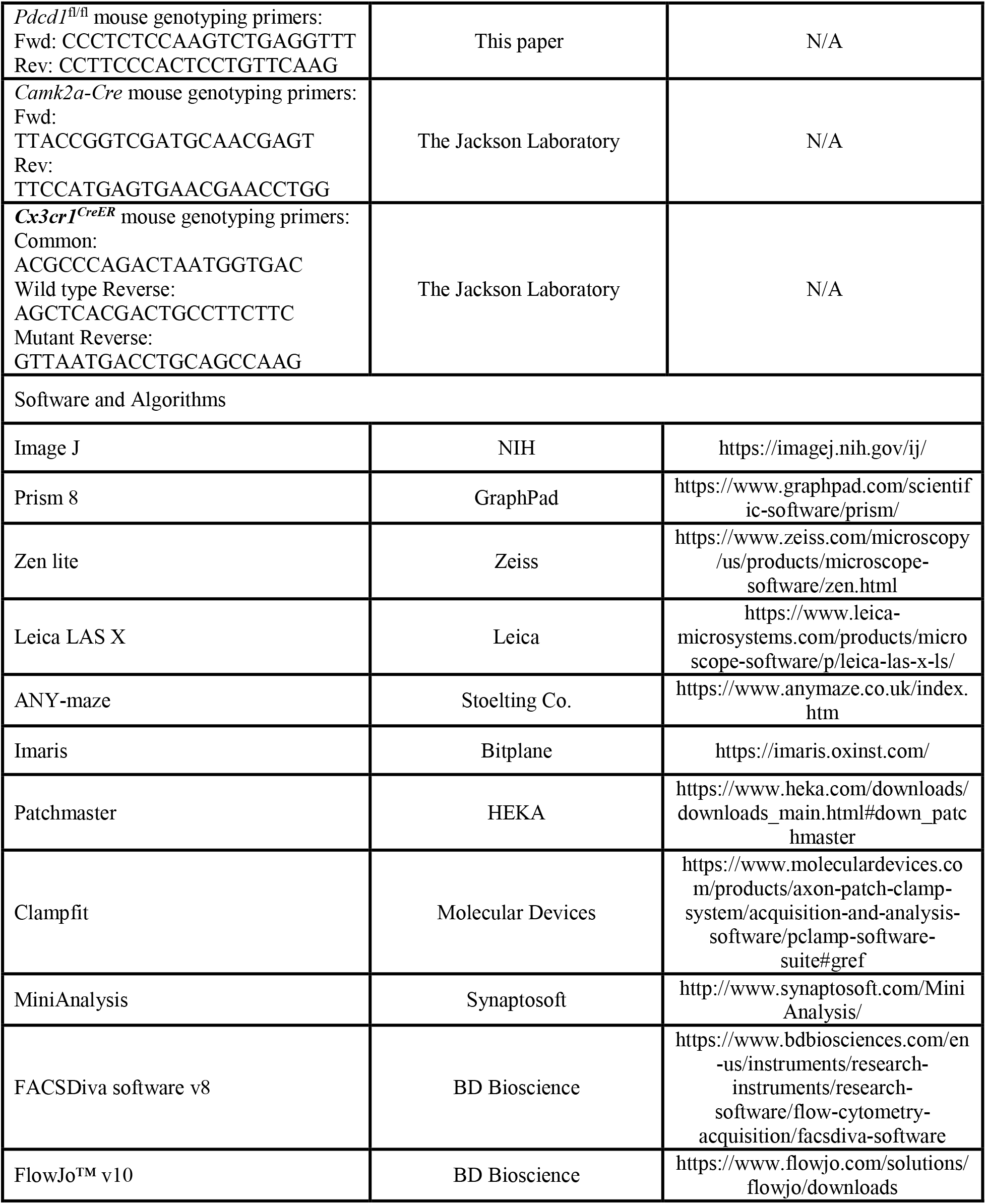

### CONTACT FOR REAGENT AND RESOURCE SHARING

Further information and requests for resources and reagents should be directed to and will be fulfilled by the lead contact, Ru-Rong Ji.

### EXPERIMENTAL MODEL AND SUBJECT DETAILS

#### Animals

Adult mice (8-16 wk old) of both sexes were used for behavioral tests, unless specifically described. Mice (5-8 weeks of both sexes) were used for electrophysiological studies. *Pdcd1* knockout mice on C57BL/6 background (Stock No:021157), *Camk2a-Cre* mice (Stock No: 005359), *Cx3cr1^CreER^* mice (Stock No: 020940) and wild-type (WT) mice (C57BL/6J, Stock No: 000664) were purchased from Jackson Laboratory. *Pdcd1^fl/fl^* mice were generated by Dr. Richard Palmiter at the University of Washington. All the mouse procedures were approved by the Institutional Animal Care & Use Committee of Duke University. All mice were maintained at Duke animal facilities and housed under a 12-hour light/dark cycle with food and water available *ad libitum*. Animals were randomly assigned to each experimental group. Two to five mice were housed in each cage. All behavioral measurements were conducted in a blinded manner and during daytime (light cycle), normally starting at 9 AM. Animal experiments were conducted in accordance with the National Institutes of Health Guide for the Care and Use of Laboratory Animals. The number of animals and sample size of each experiment were also described in Table S1. If not specified, experiments in this study were conducted in mice.

#### Generation of *Pdcd1^fl/fl^* mice and *Pdcd1* conditional knockout mice

*Pdcd1^fl/fl^* mice were generated at University of Washington. LoxP sites were engineered to flank to second exon of *Pdcd1*. The 5′ arm (∼4 kb with *Pac*I and *Sal*I sites at 5’ and 3’ ends, respectively) and 3′ arm (∼6.5 kb with *Pme*I and *Not*I sites at 5’ and 3’ ends, respectively) of the *Pdcd1* gene were PCR amplified with Q5 polymerase from a C57Bl/6 BAC clone. A loxP site was inserted into the Nhe1 site proximal to exon 2, the correct orientation verified by sequencing, and then the two arms were cloned into polylinkers of a targeting vector (4600C) that contained a loxP site, an frt-flanked Sv40-Neo gene for positive selection, HSV thymidine kinase and *Pgk*-diphtheria toxin A chain genes for negative selection (Figure S4). The construct was linearized with *Asc*1, electroporated into G4 ES cells (C57Bl/6 × 129 Sv hybrid) and correct targeting was determined by Southern blot of DNA digested with *Xba*1 using a ^32^P-labeled probe upstream of the 5’ arm of the targeting construct. Of the 62 clones analyzed 7 were correctly targeted. One clone that retained the loxP site in the 5’ arm was injected into blastocysts and resulted in good chimeras that transmitted the targeted allele through the germline. Progeny were bred with *Gt(Rosa)26Sor-FLP* recombinase mice (Jackson lab #012930) to remove the frt-flanked SV-Neo gene. Mice were then continuously backcrossed to C57Bl/6 mice. Routine genotyping is performed with 2 primers flanking the loxP site: 5′-CCCTCTCCAAGTCTGAGGTTT-3′ (*Pdcd1* F) and 5′-CCTTCCCACTCCTGTTCAAG-3′ (*Pdcd1* R). The wild-type allele gives a band of ∼150 bp, while the targeted allele with loxP site gives a band of ∼ 190 bp. Sometimes there is an additional band ∼220 bp that represents a heteroduplex (Figure S4).

*Pdcd1^fl/fl^*; *Camk2a-Cre* neuron conditional (cKO-neuron) mice were generated by crossing *Pdcd1^fl/fl^* mice with *Camk2a-*Cre mice to generate the heterozygotes *Pdcd1^fl/-^*; *Camk2a-Cre* and then crossed with *Pdcd1^fl/fl^* mice. *Pd1^fl/fl^*; *Cx3cr1^CreER^* microglia conditional (cKO-microglia) mice were generated by crossing *Pd1^fl/fl^* mice with *Cx3cr1^CreER^* mice to generate the heterozygotes *Pdcd1^fl/-^*; *Cx3cr1^CreER^* and then crossed with *Pdcd1^fl/fl^* mice. *Pd1^fl/fl^*; *Cx3cr1^CreER^* mice were injected with tamoxifen (75 mg/kg, i.p.) daily for 5 days. After the final injection, mice were kept for another 7 days before any behavioral or biochemical tests. *Pdcd1* cKO mice were validated by genotyping (Figures S5I and S5P), immunostaining (Figures S5J and S5Q) and flow cytometry (Figures 5B and 5L).

#### Mouse model of traumatic brain injury (TBI)

The closed-head TBI model (Wang et al., 2007) was established using WT or PD-1 KO mice. After anesthesia induction with 4.6% isoflurane, the trachea was intubated, and the lungs were mechanically ventilated with 1.5% isoflurane in 30% O_2_/70% N_2_. The mouse head was secured in a stereotactic device and a middle-line scalp incision was made to identify anatomical landmarks. A concave 3-mm metallic disc was adhered to the skull immediately caudal to bregma. A 2.0-mm diameter pneumatic impactor (Air-Power Inc.) was used to deliver a single midline impact to the center of the disc surface. The impactor was discharged at 6.8 ± 0.2 m/s with a head displacement of 2.5 mm. After impact, the animals could recover spontaneous ventilation before the tracheas were extubated. Mice were allowed free access to food and water after recovery. Sham mice were treated identically except for the absence of impact.

#### NHP and Human Hippocampal samples

Human hippocampus was obtained from donors through National Disease Research Interchange (NDRI) with permission of exemption from the Duke University Institutional Review Board (IRB). Postmortem whole hippocampi were dissected from 2 donors: a 59-year-old female and a 60-year-old male. NHP hippocampi were obtained from Rhesus macaques (*Macaca mulatta*) housed in the Wake Forest School of Medicine. Whole hippocampi were collected from 3 healthy monkeys: a 13-year-old female, a 19-year-old male and a 10-year-old female.

#### Primary neuronal cultures

Hippocampal neuron primary cultures were prepared from embryonic day 17-19 (E17-19) WT and PD-1 KO mice. Embryos were removed from the dams, anesthetized with isoflurane and euthanized by decapitation. Hippocampi were dissected and placed in Ca^2+^- and Mg^2+^-free Hank’s balanced salt solution (HBSS, GIBCO) and digested at 37°C in a humidified O_2_ incubator for 30 min with collagenase type II (Worthington, 285 units/mg, 12 mg/mL final concentration) and dispase II (Roche, 1 unit/mg, 20 mg/mL) in HBSS (pH 7.3). Digestion was stopped by a solution of fetal bovine serum and DMEM (GIBCO). Hippocampi were mechanically dissociated using fire-polished pipettes, filtered through a 100-µm nylon mesh and centrifuged (1,000 g, 5 min). The pellet was resuspended and plated on Poly-D-Lysine/Laminin coated glass coverslips (CORNING), and cells were plated in a DMEM solution containing 5% fetal bovine serum, 1% premixed penicillin and streptomycin. After 5-6 h, primary cultures were switched to Neurobasal Plus medium containing 2% B27 supplement, 1% GlutaMAX-I, 1% premixed penicillin and streptomycin (GIBCO). Three days after plating, cytosine arabinoside was added to a final concentration of 10 µM to curb glial proliferation. Cultures were kept at 37°C in a 5% CO_2_-humidified incubator and half of the media was replaced every three days. Whole-cell electrophysiological recording and immunocytochemistry experiments were conducted on DIV 7-8.

### METHOD DETAILS

#### Reagents

Nivolumab (OPDIVO®), a humanized anti-PD-1 antibody, was purchased from Bristol-Myers Squibb (Cat# NDC 0003-3772-11). Human IgG (Cat# ab90286) was purchased from Abcam. Probes for RNAscope were from Advanced Cell Diagnostics. All the regents for electrophysiology were purchased from Sigma or Tocris. Please find more details in STAR★METHODS and KEY RESOURCES TABLE.

#### Stereotaxic surgery and drug delivery

For stereotaxic surgery, mice were anesthetized with isoflurane (4% for induction and 1.5% thereafter for maintenance), and their heads were fixed in a stereotaxic apparatus (David Kopf Instruments). A guide cannula (62004, RWD Life Science) was stereotaxically implanted into the left lateral ventricles (AP: −0.3 mm; ML: +1.3 mm; DV: −2.0 mm) based on the mouse brain atlas (George Paxinos, 2001). After 4-day recovery, an infusion needle (62204, RWD Life Science) was inserted into lateral ventricles through the guide cannula to a depth of 2.5 mm for drug injection. Following the completion of the infusion, the needle was left for an additional 2 min to limit reflux. Control IgG or nivolumab was injected for three to six times before behavioral testing. The drug was administered at 1 µg per mouse every other day.

#### *In vivo* stereotactic viral injections

Mice were anesthetized with isoflurane (4% for induction and 1.5% thereafter for maintenance) for stereotaxic injection of viruses into the CA1 (AP: −1.8 mm, ML: ±1.4 mm, DV: −1.5 mm) and CA3 (AP: −1.8 mm; ML: ±2.2 mm; DV: −2.2 mm) for adult mice and CA1 (AP: −1.7 mm; ML: ±2.0 mm; DV: −2.1 mm) for 4-week-old mice. We injected 500 nL of virus into each location at a rate of 60 nl min^−1^ using the UltraMicroPump injection system (World Precision Instruments). The targeted hippocampal regions are shown in Figures 6A-6B and Figures S7A-7B). After each injection, the needle was left in place for an additional 10 min for efficient diffusion of the virus-containing reagent and then slowly withdrawn.

For selective re-expression of PD-1 in hippocampal neurons, PD-1 KO mice were micro-injected in the bilateral CA1 and CA3 with AAV-*Camk2a* (0.4)-*Pdcd1*-IRES-eGFP-WPRE virus (Vector Biolabs). For selective knockout of PD-1 in hippocampal neurons, *Pdcd1^fl/fl^* mice were micro-injected in the bilateral CA1 and CA3 with AAV-*Camk2a*-mCherry:Cre virus (UNC Vector Core). AAV-*Camk2a*-eGFP-WPRE virus was injected as a control. Behavioral experiments or electrophysiological recordings were performed at least 3 weeks after virus injection. Virus infection was examined at the end of all the tests.

#### Novel object recognition (NOR) test

On the first day, mice were placed in a 30 × 30 × 30 cm^3^ square arena (TAP plastics) for habituation. On the second day, two identical objects were placed in two distinct corners of the arena. Mice were then returned to their home cages for a 0.5-h or 24-h retention interval, and one of two identical objects was replaced by a new object for novel object recognition test (Zhao et al., 2017). After the retention interval, animals were placed back into the testing arena for 5 min of exploration. A valid exploration was defined as the mouse touching an object with its nose or focusing attention on the object at <1 cm. Turning around, climbing, and siting on the object was all considered to be invalid. A discrimination index was used to evaluate the scores of exploration time for the familiar or novel object (N - F/N + F) × 100%. Animal behaviors were video-recorded and analyzed by an experimenter who was blind to the testing conditions. Data were excluded if the total exploration time was less than 10 s.

#### Morris water maze (MWM) test

The Morris water maze test was conducted in a circular pool (diameter 120 cm) filled with 40-cm height water maintained at 23 ± 1°C. The water was made opaque with white milk powder. The pool is in an experimental room with many visual cues and divided into four equal quadrants. A circular platform, 8 cm in diameter, was placed in the middle of one fixed quadrant of the pool, just 1 cm underneath the water surface. During the training process, mice were trained for four trials each day on four consecutive days. Each mouse could swim for 60 s to find the platform. If the mouse failed to find the platform within the allotted time, it was picked up and placed on the platform for 5 s. The interval between trials was 20 min. In the probe test (day 5), the platform was removed and all mice were given one probe trial consisting of 60 s searching (Vorhees and Williams, 2006). If the mouse failed to cross the platform zone, the latency to platform was defined as 60 s. The swimming speed, latency to platform zone, and the number of platform zone crossings were automatically recorded and analyzed by ANY-maze (Stoelting Co.).

#### Rotarod test

A total of six trials for the rotarod test were carried out using an accelerating protocol from 4 to 60 rpm in 300 s with 20-min inter-trial intervals. Three testing sessions every day were performed for 2 days. For TBI mice, the rotarod test was measured before TBI model induction and on day1, 2, 3, 4, 5, 6, 7, 10, 14 and 21 after TBI. After falling, the mice were returned to their home cages, and the latency to fall was automatically recorded by the rotarod system (IITC Life Science Inc.). The latency was recorded as 300 s if the mice stayed on the rotarod more than 300 s.

#### Open field test

The open field test was performed in a 45 × 45 × 45 cm^3^ square arena (TAP plastics) over 15 min. The center of the arena was defined as a center area that covered 50% of the total area. The total distance travelled, and time spent in the center area were automatically recorded and analyzed by ANY-maze (Stoelting Co.).

#### Elevated plus maze test

The elevated plus maze apparatus (Stoelting Co.) is comprised of two open arms (35 × 5 cm^2^) and two closed arms (35 × 5 × 15 cm^3^) elevated 50 cm above the ground. Mice were placed in the center facing one of the two open arms and allowed to explore for 6 min. Anxiety-like behavior was assessed by time travelled within the open arms. The video and data were automatically recorded and analyzed by ANY-maze (Stoelting Co.).

#### Tail suspension test

Mice were suspended by their tails with adhesive tape (1 cm from the tail tip) and ∼15 cm away from the surface. Paper tubes were placed over the tails to ensure animals could neither climb nor hang on to their tail. The animals were video recorded for 6 min, and the time spent immobile was quantitated by an experimenter who was blind to the testing conditions. Mice were considered immobile only when they hung passively and motionlessly for at least 2 s.

#### Mouse hippocampal slices preparation

Mice were anesthetized with isoflurane and then decapitated. The brain was quickly transferred to ice-cold artificial sucrose-based cerebrospinal fluid (ACSF) containing (in mM): Sucrose 75, NaCl 87, KCl 2.5, NaH_2_PO_4_ 1.25, CaCl_2_ 0.5, MgCl_2_ 7, NaHCO_3_ 26 and glucose 25. Hippocampal slices (200-350 μm) were cut using a vibratome (Leica VT1200S). Subsequently, slices were transferred to normal ACSF solution (in mM): NaCl 124, KCl 3, NaH_2_PO_4_ 1.25, CaCl_2_ 2, MgSO_4_ 1, NaHCO_3_ 26, glucose 10 at 34°C. All extracellular solutions were constantly carbogenated (95% O_2_, 5% CO_2_). Slices were kept in ACSF for at least 1 h before recording. For PD-1 blockade expriments, slices were incubated with 300 ng/mL control IgG or nivolumab for 2 h. U0126 (1 μM, InvivoGen, Cat# tlrl-u0126) was added and incubated for 30 min before recording.

#### NHP hippocampal slice preparation

The whole hippocampus was dissected from disease-free monkeys and quickly transferred to ice-cold artificial sucrose-based cerebrospinal fluid (ACSF) containing (in mM): Sucrose 248, KCl 1, CaCl_2_ 1, MgCl_2_ 10, NaHCO_3_, 26 and glucose 10. Hippocampal slices (350 μm) were cut in ACSF with a Leica Vibratome (Leica VT1200S) within 2 hours after surgical resection (Tissues were collected at Wake Forest University and immediately transferred to Duke University for slice preparation). The slices were placed in an incubation chamber at room temperature with oxygenated recording solution for 1-5 h. The recording solution had the following composition (in mM): NaCl 124, KCl 4, CaCl_2_ 2, MgCl_2_ 2, NaHCO_3_ 26 and glucose 10. Slices were incubated with 500 ng/mL control IgG or nivolumab for 2 h before recording.

#### Whole-cell patch clamp recordings in mouse and NHP hippocampal slices and mouse hippocampal neurons of primary cultures

Whole-cell recordings were performed at 34 ± 1°C with the help of an automatic temperature controller (Warner instruments) with an EPC10 amplifier (HEKA). Data were low-pass-filtered at 2 KHz and sampled at 10 KHz. Patch pipettes were filled with a solution containing the following (in mM): K-gluconate 135, KCl 5, CaCl_2_ 0.5, MgCl_2_ 2, HEPE 5, EGTA 5, MgATP 5 (pH 7.3, 290–300 mOsm/L). When filled with the pipette solution, the resistance of the pipettes was 4-8 MΩ. RMPs, spontaneous spikes and action potentials were recorded in current-clamp mode. The action potentials were evoked by current injection steps (0 to 130 pA, 10 pA step). Data were analyzed by Patchmaster software (HEKA). For mEPSC recording, neuron was hold at −70 mV in the presence of 0.5 μM TTX and 50 μM picrotoxin to block Na^+^ currents and GABA_A_ receptors. The mEPSC were detected and analyzed using Mini Analysis (Synaptosoft Inc.). For A type current recording, the membrane voltage was held at −80 mV in the presence of 0.5 μM TTX and 2 mM CoCl_2_ to block voltage-gated Na^+^ currents and Ca^2+^ currents, and transient potassium currents (I_A_) were isolated by a two-step voltage protocol (Hu et al., 2006). Briefly, a total outward current was evoked by a command potential of +40 mV from a holding potential of −80 mV. The A-type current was dissected away from the sustained current by the voltage protocol (a 150-ms pre-pulse to −10 mV allowing the transient channels to inactivate, leaving only the sustained current). A-type current is isolated by subtraction of the sustained current from the total current. All drugs and regents for electrophysiology were purchased from Sigma or Tocris.

#### Extracellular field recordings of long-term potentiation (LTP) in brain slices

To record extracellular field excitatory postsynaptic potentials (fEPSP) in mice, a bipolar tungsten stimulating electrode (FHC) was placed along the Schaffer collateral fibers in a hippocampus slice to deliver test and conditioning stimuli, and a glass recording electrode (4-8 MΩ, filled with ACSF) was placed in the stratum radiatum of the CA1 region, 150-200 µm away from the stimulating electrode. The intensity of the stimulation was adjusted to produce a fEPSP with an amplitude of 30-40% of the maximum response. Test stimulation was delivered once per 30 s. Once a stable test response was attained, fEPSP baselines were recorded every 30 s for 10 min. Then, the LTP was induced by two HFS (100 Hz for 1 s, 30 s interval) (Shi et al., 2018). After induction, fEPSP was recorded for another 60 min. For PD-1 blockade expriments, slices were incubated with 300 ng/mL control IgG or nivolumab for 2 hours. The slope of fEPSP was analyzed by Patchmaster software (HEKA).

#### Visualization of dendrites in hippocampal neurons

To visualize the dendritic spines of mouse CA1 neurons, Neurobiotin (0.2%, Vector Laboratories) was dissolved in the intracellular pipette solution. After 30 min of whole-cell patch recording, the slices were fixed with 4% paraformaldehyde (PFA) in PBS and then processed using Alexa Fluor 594 streptavidin (1:500, Life Technologies, Cat# S32356) for visualization. Apical dendrites of CA1 neurons were imaged using a Zeiss LSM 880 with Airyscan Microscope with a 63× oil-immersion objective, and 3× optical zoom set to a Z-stack width of 0.2 mm. Imaris software (v.9.3.0, Bitplane) was used to reconstruct dendritic spines and analyze the spine density and head diameter. The dendrites were imaged from 3-4 mice for each group and analyzed similar to other studies (Tan et al., 2019; Weng et al., 2018).

#### *In situ* hybridization in mouse, NHP, and human brain sections

Mice were deeply anesthetized with isoflurane and perfused with PBS, followed by 4% PFA. Following perfusion, mouse brain was isolated and post-fixed overnight at 4°C in 4% PFA. Human brain tissues (NDRI) were delivered within 48 h of dissection and fixed in 10% formalin for >24 h. For monkey tissues, the whole hippocampus was post-fixed with 10% formalin for >24 h after dissection. After dehydration in 30% sucrose, all the tissues were embedded in OCT medium and cryosectioned to produce 14-μm tissue sections which were mounted onto the charged slides. *In situ* hybridization was performed using RNAscope® Multiplex Fluorescent Reagent Kit v2 (Advanced Cell Diagnostics, Cat# 323100) according to the manufacturer’s instructions. We used probes directed against mouse *Pdcd1* (Cat# 416781), human *PDCD1* (Cat# 602021) and customized Mmu *PDCD1*. Control probe (Cat# 700141) was used as a negative control. Following the completion of the RNAscope protocol, immunohistochemistry was performed as described in the next section. To quantify the percentages of mouse *Pdcd1* and NHP and human *PDCD1* mRNA expression in hippocampal CA1 and CA3 neurons, three randomly selected fields from CA1 and CA3 regions were analyzed from 2-3 sections of 4 mouse, 3 NHP, and 2 human samples.

#### Immunohistochemistry and imaging

Brain tissue sections (14 μm) and free-floating brain sections (30 μm) were cut in a cryostat (Leica CM1950). Tissue sections were washed several times in PBS and blocked with 0.1% Triton X-100 and 5% donkey serum for 1 h at room temperature. The sections were then incubated overnight at 4°C in a humidified chamber with the following primary antibodies: anti-PD-1 antibody (rabbit, 1:300, Sigma, Cat# PRS4065), anti-NeuN antibody (mouse, 1:1000, Millipore, Cat# MAB377), IBA1 (rabbit, 1:500, Wako, Cat# 019-19741) and nivolumab vivo 680 (1:1000, Cat# NDC 0003-3772-11). After washing, the sections were incubated with Nissl/Neuro Tracer 640/660 (1:100, Thermo Scientific, Cat# N21483) or species-specific secondary antibodies conjugated to 488-nm, 555-nm or 633-nm fluorophores (1:500, Jackson ImmunoResearch) for 2 h at room temperature. Sections were subsequently washed and coverslipped using Fluoroshield™ with DAPI (Sigma, Cat# F6057). The stained sections were examined with a Leica SP5 or Zeiss 880 confocal microscope with Z-stack and Tile Scan. The maximum projections and stitch were produced using the Zeiss Zen software. To confirm the specificity of PD-1 antibody, blocking experiments were conducted in brain sections using a mixture of anti-PD-1 antibody (1:300) and immunizing blocking peptide (1:300, Sigma, Catalog: SBP4065), based on a protocol recommended for blockade with immunizing peptide (www.abcam.com/technical). The specificity of PD-1 antibody was also tested in brain sections of PD-1 KO mice. To determine if there is neuronal loss in mice, NeuN/Nissl-positive neurons were quantified in CA1 and CA3 regions by Image J. Two or three sections were analyzed in each mouse and three or four mice per group were analyzed. All the images in the same experiment were obtained using the same settings and all analyses and quantifications were performed blinded to the experimental condition.

#### Immunocytochemistry in mouse primary hippocampal neurons

For immune staining in primary hippocampal neuron cultures (DIV 7-8) prepared from E17-19 WT and PD-1 KO embryos, neurons grown on coverslips were fixed in 4% PFA for 30 min. After washing, the coverslips with neurons were blocked with a solution containing 0.1% Triton X-100 and 5% donkey serum for 1 h at room temperature and then incubated overnight at 4°C with the following primary antibodies: anti-PD-1 antibody (rabbit, 1:300, Sigma, Cat# PRS4065) and anti-NeuN antibody (mouse, 1:1000, Millipore, Cat# MAB377). Then, the sections were incubated with species-specific secondary antibodies conjugated to 488-nm, 555-nm fluorophores secondary antibodies (1:500; Jackson ImmunoResearch) for 1 h at room temperature. Sections were subsequently washed and coverslipped using Fluoroshield™ with DAPI (Sigma, Cat# F6057). The stained sections were examined with ZEISS LSM 880 confocal microscope.

#### Flow cytometry in mouse brain tissues

Flow cytometry was conducted to characterize neurons, microglia, and immune cells in the hippocampus and cortex. It was also used to validate cKO-neuron and cKO-microglia mice. The brain tissues were isolated and digested with 1 mg/mL collagenase (Roche) for 10 min at 4°C. The digested tissue was resuspended with PBS + EDTA at 4°C and 40% Ficoll gradient (Roche) was added to remove lipid debris. To detect intracellular epitope, the cells were fixed with 4% PFA for 10 min at 4℃ and incubated with Fc receptor blocking buffer (1 µg/mL anti-mouse CD16/CD32, 2.4 G2, 2% FBS, 5% NRS, and 2% NMS 0.1 % Triton-X 100 in HBSS; BD Bioscience), and then stained with a standard panel of antibodies (see Key resource table). After staining, all cells were suspended in PBS with EDTA. Flow cytometry events were acquired using a BD FACS Canto II flow cytometer using the FACSDiva software v8 software (BD Bioscience). The FCS files were analyzed using FlowJo™ v10 (BD bioscience).

#### Antibody labeling with dye tracer

To visualize the antibody binding to tissue sections, nivolumab (1 mg) and control IgG were conjugated with Vivo-680 (10 μg) dye in 0.1 mol/L sodium bicarbonate buffer overnight at 4°C, according to the manufacturer’s instructions (PerkinElmer, Cat# NEV11118). CamKIIα and control IgG antibodies were conjugated with Alexa flour 488 dye (Invitrogen, Cat# A10468) in 0.1 mol/L sodium bicarbonate buffer overnight at 4℃. Specific antibody labeling was predicated with a gel-filtration filter and then dilated with PBS. The salt and non-binding dye were removed by dialysis (4°C, 24 h) using a 10 kDa cut off membrane (Sigma). The labeled antibodies were calculated by Nano-drop.

#### ELISA

Mouse TNF-α and IL-17 ELISA kits were purchased from the R&D system (Cat# DY410 for TNF-α and Cat# DY421 for IL-17). Mouse IL-1β ELISA kit was purchased from the Biolegend (Cat# 432601).The brain tissues were digested with RIPA buffer (Sigma) containing protease inhibitor cocktail (Sigma). ELISA assays were conducted according to the manufacturer’s instructions. 100 μg of proteins were used for each reaction. The absorbance was measured in 450 nm wavelength using a 96-well xMark™ Microplate plate reader (BioRad). The cytokine levels were calculated by standard curve and analyzed using microplate manager 6 software (BioRad). The tissue cytokine level was normalized by total protein level which was calculated using BCA assay. Standard curve was included for each measurement.

### QUANTIFICATION AND STATISTICAL ANALYSIS

All data were expressed as the mean ± SEM. The sample size and statistical analysis for each experiment were indicated in the figures and figure legends. No statistical method was used to predetermine sample size. Sample sizes were estimated based on our previous studies for similar types of behavioral, biochemical, and electrophysiological analyses (Berta et al., 2014; Chen et al., 2015; Xu et al., 2015). Statistical analyses were completed with Prism GraphPad 8.0. The criterion for statistical significance was *P* < 0.05. The detailed statistical analyses also described in Table S2.

### DATA AND SOFTWARE AVAILABILITY

Original data are available upon request.

No custom software was used in this study.

---

**Figure S1.**
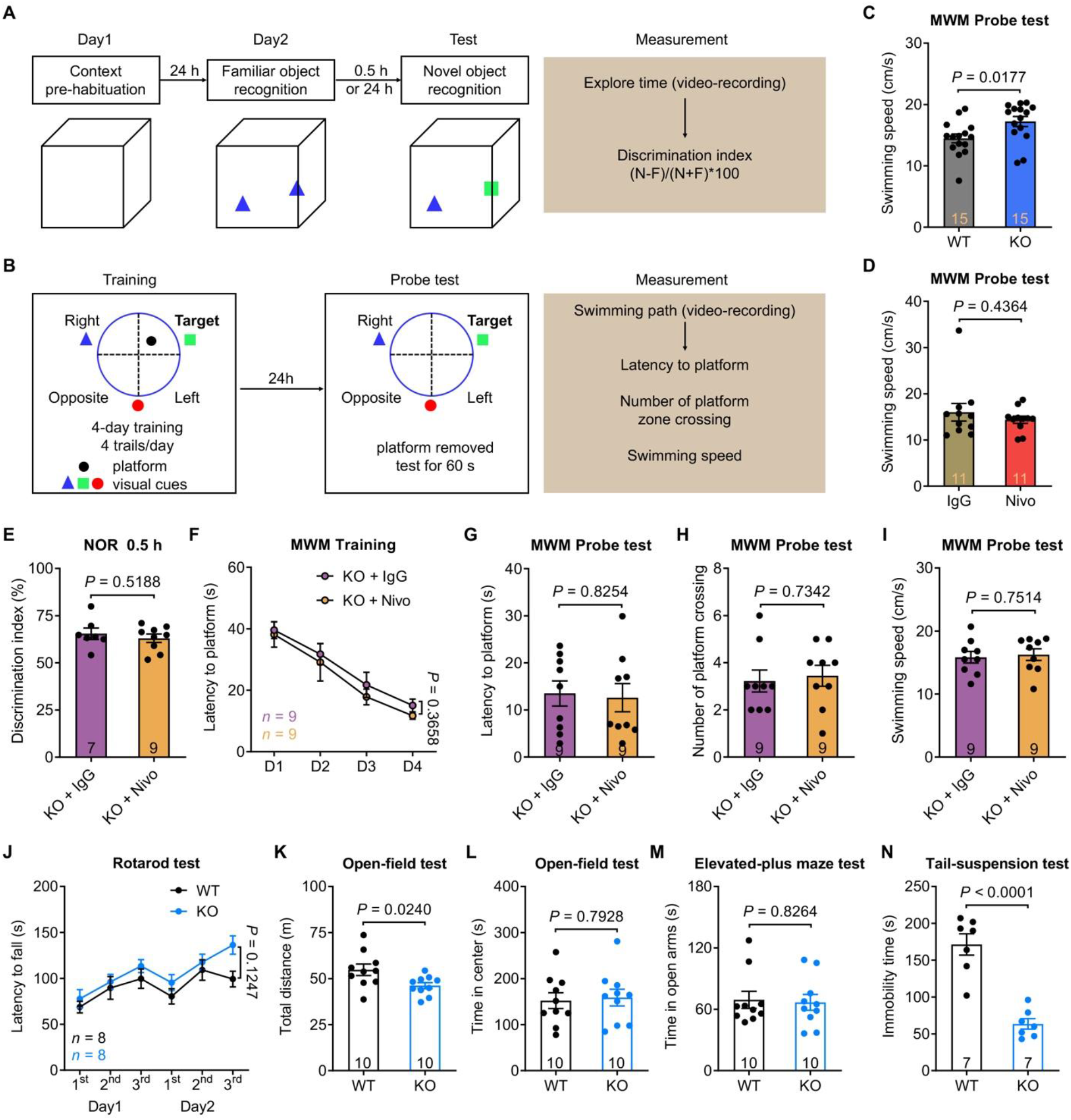
Behavioral tests for novel object recognition (NOR), Morris water maze (MWM) and anxiety/depression in WT and PD-1 KO mice. Related to Figure 1. **(A)** Schematic of NOR testing. **(B)** Schematic of MWM training and probe tests. **(C)** MWM testing shows higher swimming speed in PD-1 KO vs. WT mice (n = 15, *P* = 0.0177). **(D)** MWM probe test shows comparable swimming speed (*P* = 0.4364) in WT mice treated with control IgG (n = 11) and nivolumab (n = 11). **(E-I)** ICV nivolumab treatment has no effects on cognition in PD-1 KO mice. **(E)** NOR testing shows comparable learning capacity in PD-1 KO mice treated with control IgG (n = 7) and nivolumab (n = 9, *P* = 0.5188). Control IgG and nivolumab (3 × 1 μg) was ICV administrated every other day. **(F)** MWM training test shows no difference (*P* = 0.3658) in PD-1 KO mice treated with control IgG (n = 9) and nivolumab (n = 9). Control IgG or nivolumab (5 × 1 μg) was ICV administrated every other day (F-I). **(G-I)** MWM probe tests show no difference in latency to platform (G, *P* = 0.8254), platform crossing number (H, *P* = 0.7342) and swimming speed (I, *P* = 0.7514) in PD-1 KO mice treated with control IgG (n = 9) and nivolumab (n = 9). **(J)** Motor coordination is comparable in WT mice (n = 8) and PD-1 KO mice (n = 8) in rotarod test (*P* = 0.1247). **(K)** Locomotor activities of WT mice (n = 10) and PD-1 KO mice (n = 10) in open-field test (*P* = 0.0240). **(L-M)** Anxiety-like behaviors of WT mice (n = 10) and PD-1 KO mice (n = 10) measured by open-field test (L, *P* = 0.7928) and elevated-plus maze test (M, *P* = 0.8264). **(N)** Depressive-like behaviors of WT mice (n = 7) and PD-1 KO mice (n = 7) measured by tail-suspension test. Note that PD-1 KO mice show significant improvement in depression (*P* < 0.0001). Data are represented as mean ± SEM. Two-tailed Student’s t test (C, D, E, G, H, I, K, L, M, N), Two-way ANOVA (F, J).

**Figure S2.**
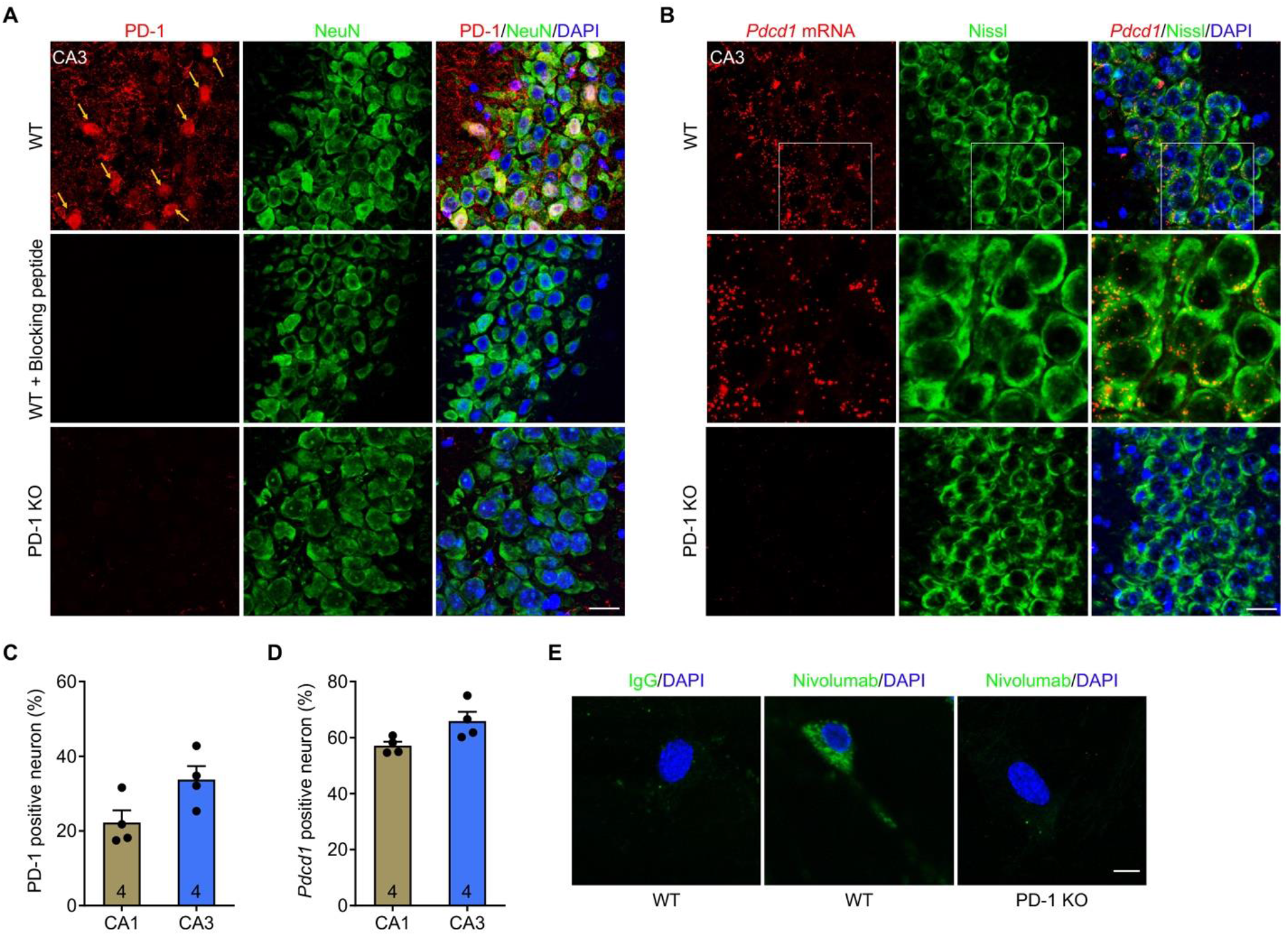
Characterization of PD-1 and *Pdcd1* mRNA expression in mouse hippocampal neurons. Related to Figure 2. **(A)** Representative images of PD-1 immunostaining in WT CA3 neurons, which is lost after incubation with PD-1 blocking peptide or in PD-1 KO neurons. Arrows indicate PD-1 positive neurons. Scale bar, 20 µm. **(B)** Representative images of *Pdcd1 in situ* hybridization showing *Pdcd1* mRNA expression of PD-1 in WT but not in PD-1 KO CA3 neurons. Higher magnification images (middle) show co-localization of *Pdcd1* and Nissl. Scale bar, 20 µm. **(C)** Quantification of PD-1 positive neurons in WT CA1 and CA3 neurons. n = 4 mice. **(D)** Quantification of *Pdcd1* mRNA positive neurons in WT CA1 and CA3 neurons. n = 4 mice. **(E)** Representative images of immunocytochemistry show nivolumab binding on primary hippocampal neurons from WT but not in PD-1 KO mice. Scale bar, 10 µm. Data are represented as mean ± SEM.

**Figure S3.**
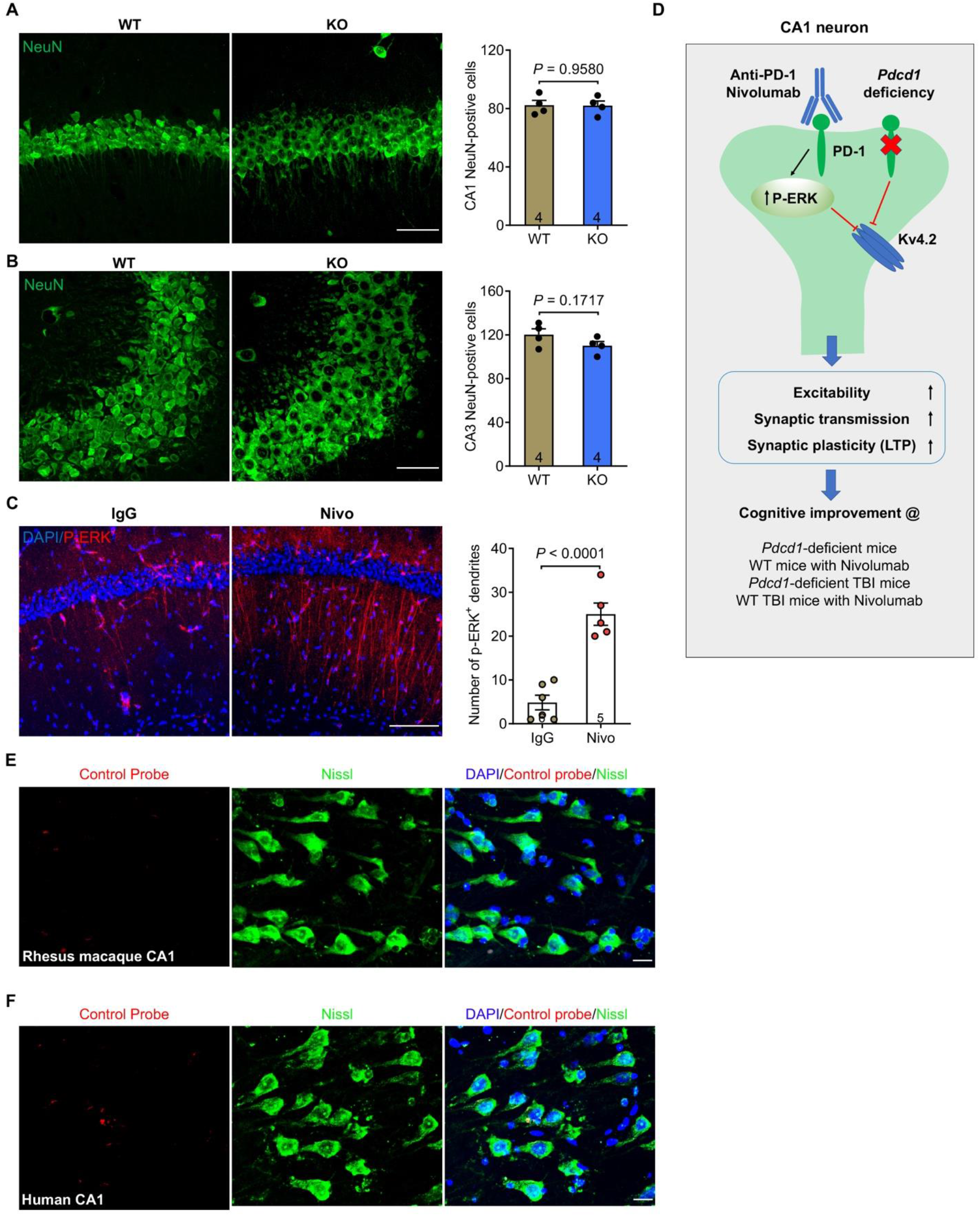
Quantification of hippocampal neurons in WT and PD-1 KO mice, induction of P-ERK by Nivolumab, and working hypothesis. Related to Figure 3 and Figure 4. **(A-B)** Confocal microscopy images and quantification of NeuN-positive neurons in CA1 (A) and CA3 (B) of WT (n = 4) and PD-1 KO mice (n = 4). Note PD-1 KO and WT mice have no differences in neuron numbers in CA1 (*P* = 0.9580) and CA3 (*P* = 0.1717). Scale bar, 50 µm. **(C)** Confocal microscopy images P-ERK immunostaining (left) and quantification of the number of P-ERK-positive dendrites (right) in CA1 of brain slices incubated with control IgG (n = 6 slices from 3 mice) and nivolumab (n = 5 slices from 3 mice, 300 ng/mL for 2 h). Note that nivolumab treatment significantly increases number of P-ERK^+^ CA1 dendrites (*P* < 0.0001). Scale bar, 100 µm. **(D)** Schematic illustration of working hypothesis by which PD-1 regulates synaptic function and cognition in CA1 neurons. Loss of PD-1 in PD-1 KO mice or blockade of PD-1 with Nivolumab in WT mice enhances cognition in both physiological and pathological (TBI) conditions. Loss of PD-1 or blockade of PD-1 also results in increases in neuronal excitability, synaptic transmission, and LTP, as well as a reduction of Kv4.2-mediated A-type K^+^ currents in hippocampal CA1 neurons. Nivolumab induces P-ERK in CA1 dendrites to suppress A-type K^+^ currents, leading to changes in neuronal excitability and synaptic plasticity in learning and memory. **(E-F)** Representative images of in situ hybridization show no staining of negative control probe in hippocampal neurons of Rhesus macaque (E) and human (F) brain sections. Scale bar, 20 µm. Data are represented as mean ± SEM. Two-tailed Student’s t test (A, B, C).

**Figure S4.**
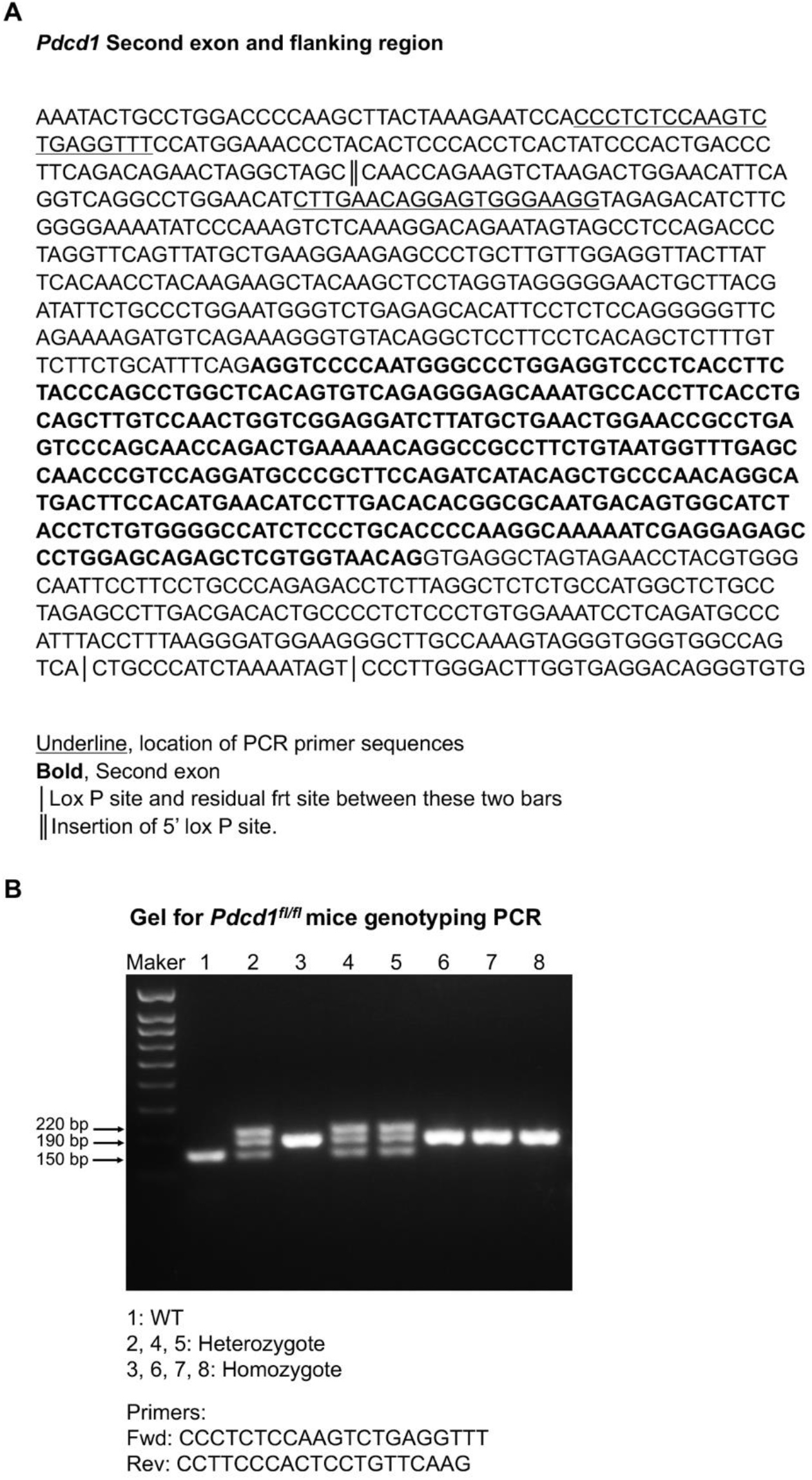
Schematic of generating *Pdcd1^fl/fl^* mice and validation of transgenic mice. **(A)** *Pdcd1* sequence including second exon (bold), Lox P site and residual frt site, and Insertion of 5’ lox P site. The sequences for the design of PCR primers are underlined. (B) Validation of *Pdcd1^fl/fl^* mice *via* genotyping. PCR primer sequences are included.

**Figure S5.**
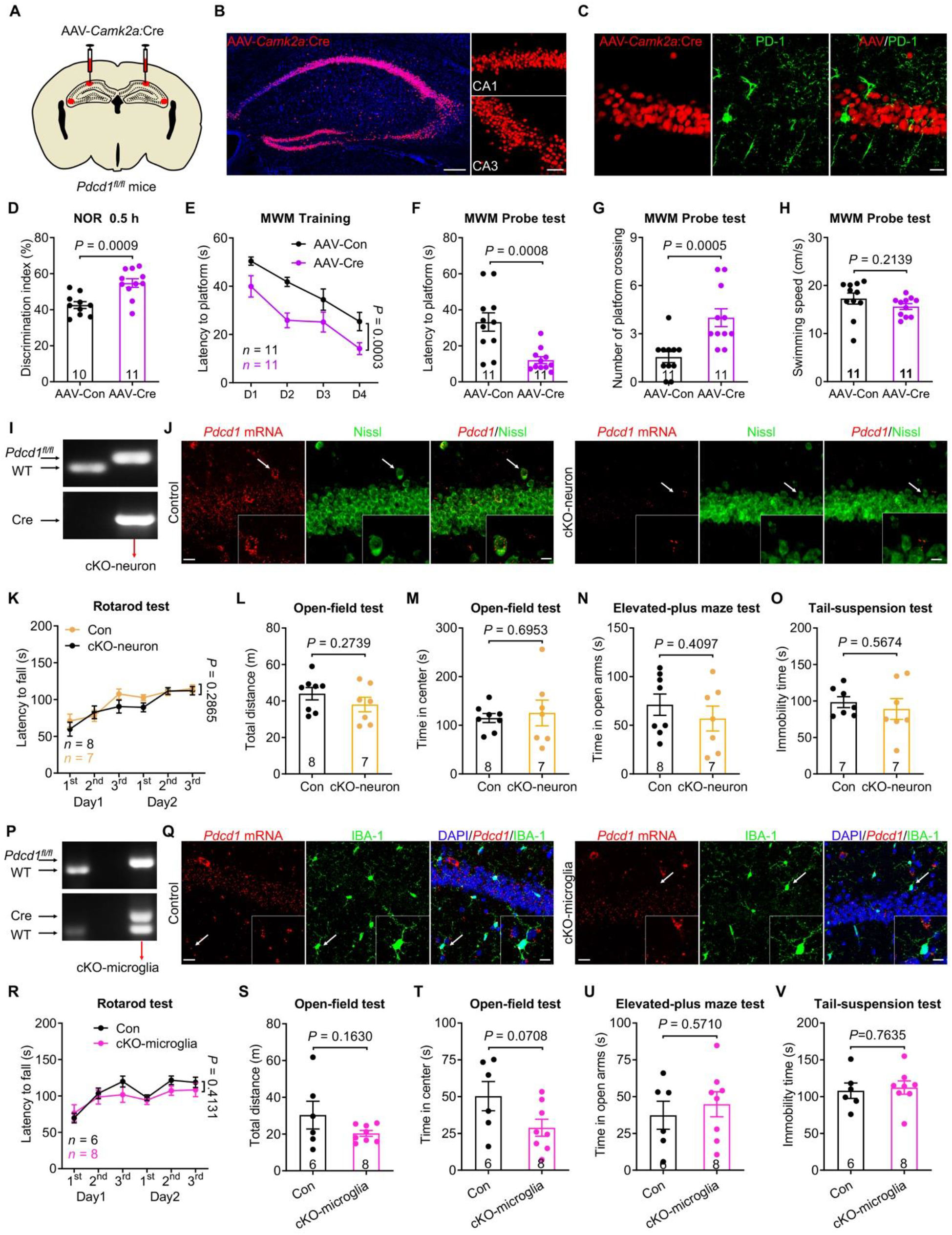
Selective deletion of *Pdcd1* in hippocampal excitatory neurons but not microglia enhances the learning and memory. Related to Figure 5. **(A)** Illustration of bilateral virus (AAV-Camk2a-Cre or AAV-control) injections into hippocampus of *Pdcd1^fl/fl^* mice to selectively delete *Pdcd1* in hippocampal excitatory neurons. **(B)** Representative confocal images of the mouse hippocampus show the AAV infected neurons. Blue: DAPI. Scale bar, 200 µm. Right, higher magnification images show infected CA1 and CA3 excitatory neurons. Scale bar, 50 µm. **(C)** Representative confocal images show the selective loss of PD-1 in CA1 excitatory neurons that are AAV-CaMKIIα-Cre positive. Scale bar, 20 µm. **(D)** NOR test in *Pdcd1^fl/fl^* mice injected with AAV-control (n = 10) and AAV-Camk2a-Cre (n = 11). Note that *Pdcd1^fl/fl^* mice injected with AAV-CaMKIIα-Cre exhibit significant improvement (*P* = 0.0009). **(E)** Spatial learning curves during MWM training in *Pdcd1^fl/fl^* mice show better performance in mice injected with AAV-Camk2a-Cre vs. mice injected with AAV-control (n = 11, *P* = 0.0003). **(F-G)** MWM probe tests show stronger performance in *Pdcd1^fl/fl^* mice injected with AAV-Camk2a-Cre vs. AAV-control (n = 11) in latency to platform (F, *P* = 0.0008) and platform zone crossings number (G, *P* = 0.0005). **(H)** MWM probe testing shows the comparable swimming speed in *Pdcd1^fl/fl^* mice injected with AAV-control vs. AAV-Camk2a-Cre (n = 11, *P* = 0.2139). **(I)** Genotyping characterization of cKO-neuron mice. **(J)** Representative images show *Pdcd1* expression in CA1 neurons of control littermates, which is markedly reduced in cKO-neuron mice. Scale bar, 20 µm. Inserted boxes are higher magnification images showing co-localization of *Pdcd1* with Nissl in CA1 neurons. Scale bar, 10 µm. Arrows indicate the enlarged neuron in the insert. **(K)** Motor coordination is comparable in control littermates (n = 8) and cKO-neuron mice (n = 7) in rotarod test (*P* = 0.2865). **(L)** Locomotor activities are unaltered in control littermates (n = 8) and cKO-neuron mice (n = 7) in open-field test (*P* = 0.2739). **(M-N)** Anxiety-like behaviors are comparable in control littermates (n = 8) and cKO-neuron mice (n = 7) in open-field test (*P* = 0.6953) and elevated-plus maze test (*P* = 0.4097). **(O)** Depressive-like behaviors are comparable in control littermates (n = 7) and cKO-neuron mice (n = 7) in tail-suspension test (*P* = 0.5674). **(P)** Genotyping characterization of cKO-microglia mice. **(Q)** Representative images show *Pdcd1* is expression in CA1 microglia of control littermates, which is substantially reduced in cKO-microglia mice. Inserted boxes are higher magnification images showing co-localization of *Pdcd1* with IBA-1 in CA1 neurons. Scale bar, 10 µm. Arrows indicate the enlarged microglial cell in the insert. **(R)** Motor coordination is comparable in control littermates (n = 6) vs. cKO-microglia mice (n = 8) in rotarod test (*P* = 0.4131). **(S)** Locomotor activities are comparable in control littermates (n = 6) vs. cKO-microglia mice (n = 8) in open-field test (*P* = 0.1630). **(T-U)** Anxiety-like behaviors are not significantly altered in cKO-microglia mice (n = 8) compared to control littermates (n = 6) in open-field test (T, *P* = 0.0708) and elevated-plus maze test (U, *P* = 0.5710). **(V)** Depressive-like behavior is comparable in control littermates (n = 6) and cKO-microglia mice (n = 8) measured by tail-suspension test (*P* = 0.7635). Data are represented as mean ± SEM. Two-tailed Student’s t test (D, H, L, M, N, O, S, T, U, V), Two-way ANOVA (E, K, R), Mann-Whitney test (F, G).

**Figure S6.**
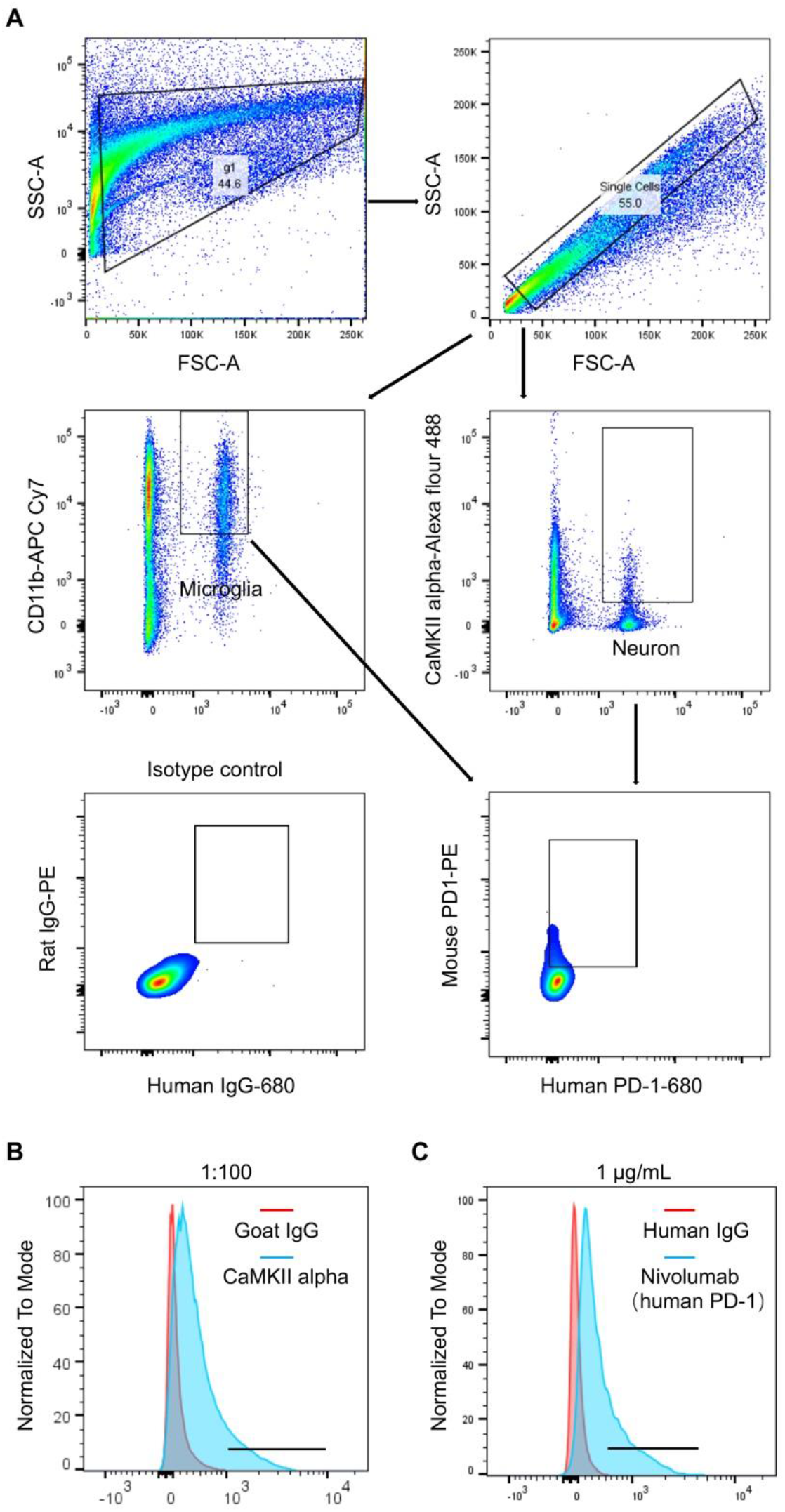
Flow cytometry analysis of conditional knockout mice. Related to Figure 5. **(A)** Flow cytometry gating strategy for testing PD-1 expression in neurons and microglia and the validation cKO-neuron mice and cKO-microglia mice. **(B,C)** Validation of florescence-labeled CaMKII alpha-Alexa flour 488 antibody (B) and Nivolumab (human PD-1)-vivo 680 antibody (C).

**Figure S7.**
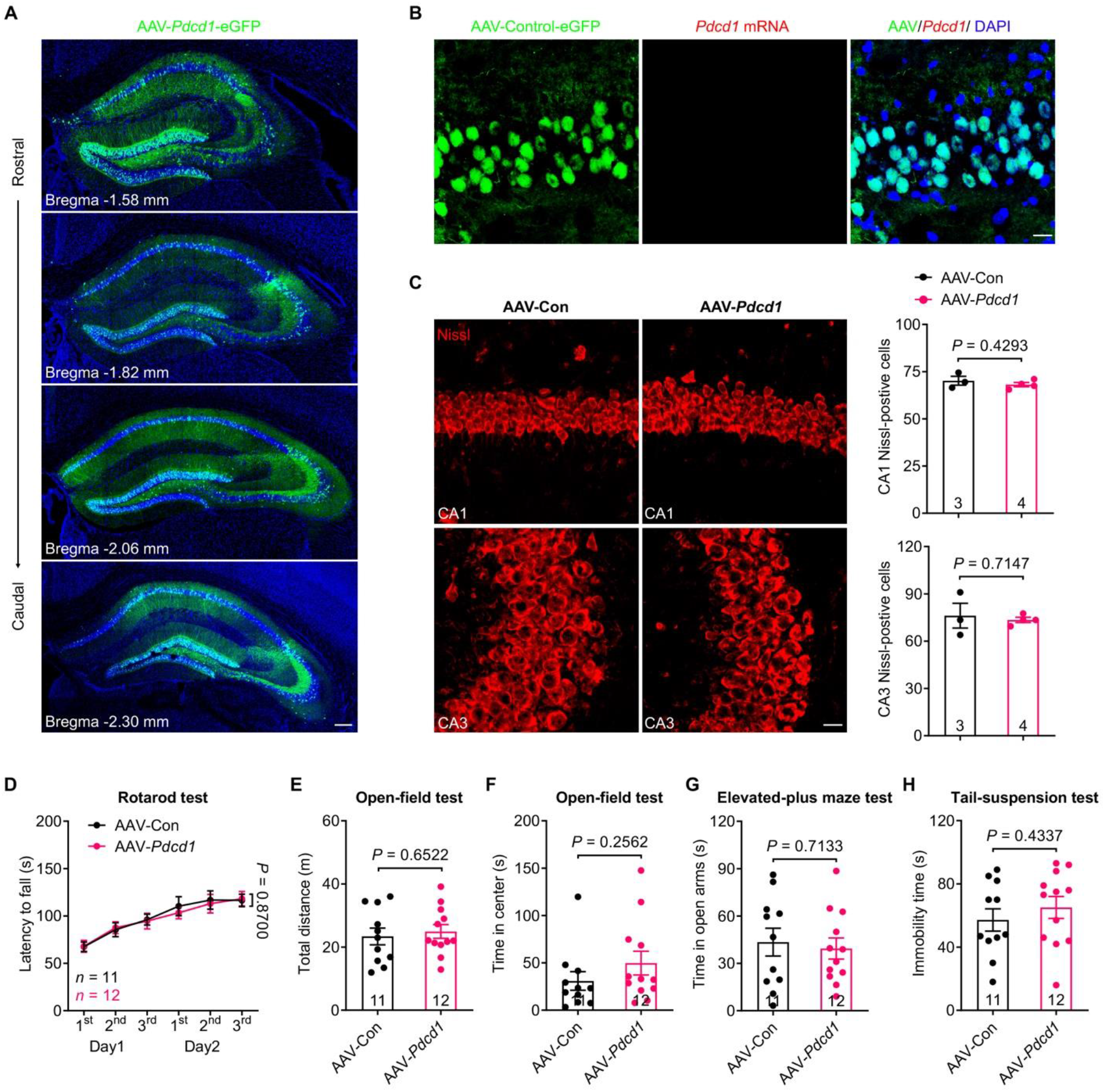
Characterization of *Pdcd1* expression, neuronal number, and behaviors in PD-1 KO mice with the AAV-*Pdcd1* injections into the hippocampus. Related to Figure 6. **(A)** Representative confocal images of brain sections (rostral to caudal orientation) from the mice injected with AAV-*Pdcd1* virus. Blue: DAPI. Scale bar, 200 µm. **(B)** Representative confocal images show no *Pdcd1* re-expression in PD-1 KO mice injected with AAV-control virus. Scale bar, 20 µm. **(C)** Representative confocal images and quantification of CA1 and CA3 neurons in PD-1 KO mice injected with AAV-control (n = 3) and AAV-*Pdcd1* (n = 4). AAV-*Pdcd1* treatment does not change neuron number in CA1 (*P* = 0.4293) and CA3 (*P* = 0.7147). Scale bar, 20 µm. **(D)** Motor coordination is comparable (*P* = 0.8700) in *Pd1*^−/−^ mice injected with AAV-control (n = 11) and AAV-*Pdcd1* (n = 12) in rotarod test. **(E)** Locomotor function is comparable (*P* = 0.6522) in PD-1 KO mice injected with AAV-control (n = 11) and AAV-*Pdcd1* (n = 12) in open-field test. **(F-G)** Anxiety-like behaviors are comparable in *Pd1*^−/−^ mice injected with AAV-control (n = 11) and AAV-*Pdcd1* (n = 12) measured by open-field test (F, *P* = 0.2562) and elevated-plus maze test (G, *P* = 0.7133). **(H)** Depressive-like behavior is comparable (*P* = 0.4337) in PD-1 KO mice injected with AAV-control (n = 11) and AAV-*Pdcd1* (n = 12) measured by tail-suspension test. Two-tailed Student’s t test, n: the number of mice. Data are represented as mean ± SEM. Two-tailed Student’s t test (C, E, F, G, H), Two-way ANOVA (D).

**Figure S8.**
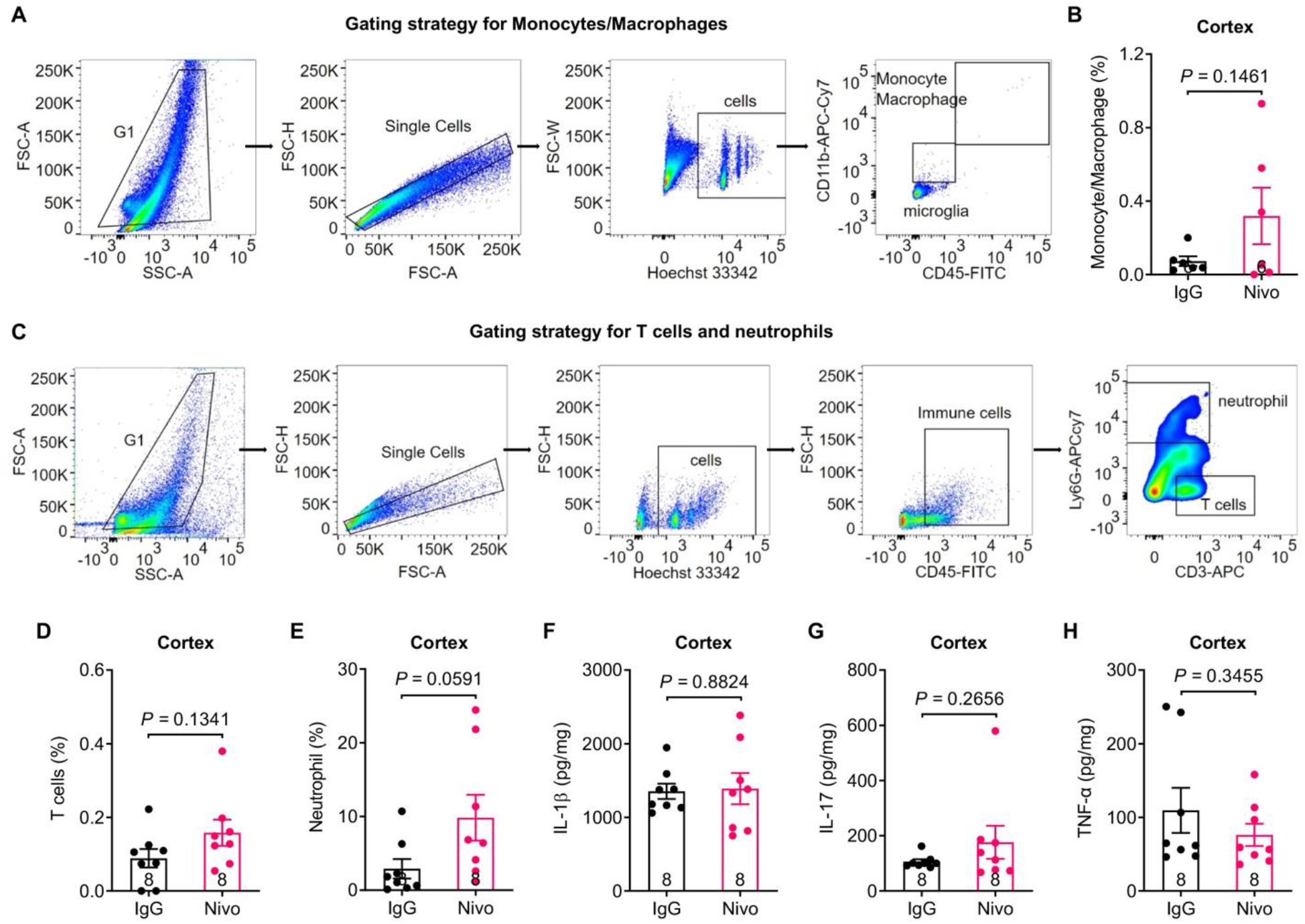
Flow cytometry and ELISA analyses in TBI mice treated with control IgG and nivolumab. Related to Figure 7. **(A)** Flow cytometry gating strategy for monocytes/macrophages. **(B)** Flow cytometry of cortex tissues from control IgG and nivolumab (n = 6) treated mice analyzed for the percentage of monocytes/macrophages (*P* = 0.1461). **(C)** Flow cytometry of gating strategy for T cells and neutrophils. **(D-E)** Flow cytometry of cortex tissues from control IgG and nivolumab (n = 8) treated mice analyzed for the percentage of T cells (D, *P* = 0.1341) and neutrophil (E, *P* = 0.0591). **(F-H)** ELISA test of cortex tissues from control IgG and nivolumab treated mice (n = 8) analyzed for the levels of pro-inflammatory cytokines (F: IL-1β *P* = 0.8824; G: IL-17 *P* = 0.2656; H: TNF-α *P* = 0.3455). Data are represented as mean ± SEM. Two-tailed Student’s t test (B, D, E, F, G, H).

**Table S1.** Number of mice, NHPs and human donors used in this study. Related to Figures 1-7 and Figures S1-8. (note: mice shared in different experiments are underlined.)

| Figures | Sample size |  | Number of groups | Number of samples | Sex | Ages | Total number |
| --- | --- | --- | --- | --- | --- | --- | --- |
| Figure 1A | 9, 6 | mice | 2 | 15 | Male | 8-12 weeks | 15 |
| Figure 1B | 9, 9 | mice | 2 | 18 | Male | 8-12 weeks | 18 |
| Figure 1C-E<br>Figure S1C | 15, 15 | mice | 2 | 30 | Male | 8-12 weeks | 30 |
| Figure 1F | 9, 8 | mice | 2 | 17 | Male | 8-12 weeks | 17 |
| Figure 1G | 6, 8 | mice | 2 | 14 | Male | 8-12 weeks | 14 |
| Figure 1H-J<br>Figure S1D | 11, 11 | mice | 2 | 22 | Male | 8-12 weeks | 22 |
| Figure 1K | 10, 7 | mice | 2 | 17 | 5 male, 5 female<br>4 male, 3 female | 40 weeks | <u>17</u> |
| Figure 1L-O | 13, 10 | mice | 2 | 23 | 9 male, 4 female<br>6 male, 4 female | 40 weeks | 23 |
| Figure S1E | 7, 9 | mice | 2 | 16 | male | 8-12 weeks | <u>16</u> |
| Figure S1F-I | 9, 9 | mice | 2 | 18 | male | 8-12 weeks | 18 |
| Figure 1J | 8, 8 | mice | 2 | 16 | 4 male, 4 female<br>4 male, 4 female | 8-12 weeks | 16 |
| Figure 1K-L | 10, 10 | mice | 2 | 20 | 5 male, 5 female<br>5 male, 5 female | 8-12 weeks | 20 |
| Figure 1M | 10, 10 | mice | 2 | 20 | 5 male, 5 female<br>5 male, 5 female | 8-12 weeks | 20 |
| Figure 1N | 7, 7 | mice | 2 | 14 | male | 8-12 weeks | 14 |
| Figure 2D-F | 20/6,<br>20/3 | neurons/<br>mice | 2 | 40 | 3 male, 3 female<br>2 male, 1 female | 5-7 weeks | 9 |
| Figure 2G-H | 16/3,<br>16/4 | neurons/<br>mice | 2 | 32 | 2 male, 1 female<br>2 male, 2 female | 5-7 weeks | 7 |
| Figure 2K | 16/18,<br>18/10 | neurons/<br>embryos | 2 | 34 |  | E17-19 | 28 |
| Figure 2L | 16/18,<br>13/10 | neurons/<br>embryos | 2 | 29 |  | E17-19 | <u>28</u> |
| Figure S2C | 4, 4 | mice | 2 | 8 | 2 male, 2 female | 8-12 weeks | 4 |
| Figure S2D | 4, 4 | mice | 2 | 8 | 2 male, 2 female | 8-12 weeks | 4 |
| Figure 3A | 13/3,<br>13/4 | neurons/<br>mice | 2 | 26 | 2 male, 1 female<br>2 male, 2 female | 5-7 weeks | 7 |
| Figure 3B | 14/4,<br>14/3 | neurons/<br>mice | 2 | 28 | 2 male, 2 female<br>2 male, 1 female | 5-7 weeks | 7 |
| Figure 3C-D | 5/5,<br>5/5 | slices/<br>mice | 2 | 10 | 3 male, 2 female<br>4 male, 1 female | 6-8 weeks | 10 |
| Figure 3E-F | 5/5,<br>6/6 | slices/<br>mice | 2 | 11 | 3 male, 2 female<br>3 male, 3 female | 6-8 weeks | 11 |
| Figure 3H | 33/4,<br>34/4 | dendrites<br>/mice | 2 | 67 | 3 male, 1 female<br>3 male, 1 female | 6-8 weeks | 8 |
| Figure 3I | 33/4<br>34/4 | dendrites<br>/mice | 2 | 4186 | 3 male, 1 female<br>3 male, 1 female | 6-8 weeks | <u>8</u> |
| Figure 3K | 34/3,<br>42/3 | dendrites<br>/mice | 2 | 76 | 2 male, 1 female<br>1 male, 2 female | 6-8 weeks | 6 |
| Figure 3L | 34/3<br>42/3 | dendrites<br>/mice | 2 | 4594 | 2 male, 1 female<br>1 male, 2 female | 6-8 weeks | <u>6</u> |
| Figure 3M | 17/3<br>18/3<br>20/3<br>15/3 | neurons/<br>mice | 4 | 70 | 2 male, 1 female<br>2 male, 1 female<br>1 male, 2 female<br>2 male, 1 female | 5-7 weeks | 12 |
| Figure 3N | 15/3<br>14/3<br>12/3<br>11/3 | neurons/<br>mice | 4 | 52 | 1 male, 2 female<br>2 male, 1 female<br>1 male, 2 female<br>2 male, 1 female | 5-7 weeks | 12 |
| Figure S3<br>A-B | 4, 4 | mice | 2 | 8 | 2 male, 2 female<br>2 male, 2 female | 8-12 weeks | 8 |
| Figure S3C | 6/3,<br>5/3 | slices/<br>mice | 2 | 11 | 2 male, 1 female<br>3 male | 6-8 weeks | 6 |
| Figure 4B | 3, 3 | NHPs | 2 | 6 | 1 male, 2 female | 10-19 years | 3 |
| Figure 4D | 19/3,<br>18/3 | neuron/<br>NHPs | 2 | 37 | 1 male, 2 female | 10-19 years | <u>3</u> |
| Figure 4F | 2, 2 | human | 2 | 4 | 1 male, 1 female | 59-60 years | 2 |
| Figure 5B | 6, 5, 6 | mice | 3 | 17 | 3 male, 3 female<br>3 male, 2 female<br>3 male, 3 female | 8-12 weeks | 17 |
| Figure 5C | 8, 6 | mice | 2 | 14 | 5 male, 3 female<br>4 male, 2 female | 8-12 weeks | <u>14</u> |
| Figure 5D-G | 8, 7 | mice | 2 | 15 | 5 male, 3 female<br>5 male, 2 female | 8-12 weeks | 15 |
| Figure 5I-J | 6/5<br>5/5 | slices/<br>mice | 2 | 11 | 3 male, 2 female<br>3 male, 2 female | 6-8 weeks | 10 |
| Figure 5L | 6, 6, 5 | mice | 3 | 17 | 3 male, 3 female<br>3 male, 2 female<br>3 male, 3 female | 8-12 weeks | <u>17</u> |
| Figure 5M | 11, 10 | mice | 2 | 21 | 9 male, 2 female<br>8 male, 2 female | 8-12 weeks | 21 |
| Figure 5N-Q | 11, 10 | mice | 2 | 21 | 9 male, 2 female<br>8 male, 2 female | 8-12 weeks | <u>21</u> |
| Figure S5D | 10, 11 | mice | 2 | 21 | 8 male, 2 female<br>9 male, 2 female | 8-12 weeks | <u>21</u> |
| Figure S5<br>E-H | 11, 11 | mice | 2 | 22 | 9 male, 2 female<br>9 male, 2 female | 8-12 weeks | 22 |
| Figure S5<br>K-O | 8, 7 | mice | 2 | 15 | 6 male, 2 female<br>5 male, 2 female | 8-12 weeks | <u>15</u> |
| Figure S5<br>R-V | 6, 8 | mice | 2 | 14 | male | 8-12 weeks | <u>14</u> |
| Figure 6E | 11, 11 | mice | 2 | 22 | 9 male, 2 female<br>9 male, 2 female | 8-12 weeks | <u>22</u> |
| Figure 6F-I | 11, 12 | mice | 2 | 23 | 9 male, 2 female<br>10 male, 2<br>female | 8-12 weeks | 23 |
| Figure 6K-L | 6/5,<br>6/5 | mice | 2 | 12 | 3 male, 2 female<br>4 male, 1 female | 8-10 weeks | 10 |
| Figure S7C | 3, 4 | mice | 2 | 7 | 2 male, 1 female<br>2 male, 2 female | 8-12 weeks | <u>7</u> |
| Figure S7<br>D-H | 11, 12 | mice | 2 | 23 | 9 male, 2 female<br>10 male, 2<br>female | 8-12 weeks | <u>23</u> |
| Figure 7<br>A-F | 8, 7, 9,<br>8 | mice | 4 | 32 | male | 8-12 weeks | 32 |
| Figure 7<br>G-R | 8, 8 | mice | 2 | 16 | male | 8-12 weeks | 16 |
| Figure S8B | 6, 6 | mice | 2 | 12 | male | 8-12 weeks | <u>12</u> |
| Figure S8<br>D-H | 8, 8 | mice | 2 | 16 | male | 8-12 weeks | <u>16</u> |
| Total<br>number |  |  |  |  |  |  | 532 mice<br>3 NHPs<br>2 donors |

**Table S2.** Statistical information. Related to Figures 1-7 and Figures S1-8.

| Figure | Statistical method | Statistical result |
| --- | --- | --- |
| Figure 1A | Two-tailed Student's t test | $t = 2.419, P = 0.0309$ |
| Figure 1B | Two-tailed Student's t test | $t = 2.406, P = 0.0286$ |
| Figure 1C | Two-way ANOVA | $F(1, 28) = 17.35, P = 0.0003$ |
| Figure 1D | Mann-Whitney test | $U = 25, P = 0.0001$ |
| Figure 1E | Mann-Whitney test | $U = 41, P = 0.0019$ |
| Figure 1F | Two-tailed Student's t test | $t = 3.324, P = 0.0046$ |
| Figure 1G | Two-tailed Student's t test | $t = 3.729, P = 0.0029$ |
| Figure 1H | Two-way ANOVA | $F(1, 20) = 6.459, P = 0.0194$ |
| Figure 1I | Mann-Whitney test | $U = 29, P = 0.0398$ |
| Figure 1J | Mann-Whitney test | $U = 29, P = 0.0344$ |
| Figure 1K | Two-tailed Student's t test | $t = 3.761, P = 0.0019$ |
| Figure 1L | Two-way ANOVA | $F(1, 21) = 11.42, P = 0.0028$ |
| Figure 1M | Mann-Whitney test | $U = 4, P < 0.0001$ |
| Figure 1N | Mann-Whitney test | $U = 7.5, P < 0.0001$ |
| Figure 1O | Two-tailed Student's t test | $t = 1.553, P = 0.1353$ |
| Figure S1C | Two-tailed Student's t test | $t = 2.520, P = 0.0177$ |
| Figure S1D | Two-tailed Student's t test | $t = 0.7943, P = 0.4364$ |
| Figure S1E | Two-tailed Student's t test | $t = 0.6619, P = 0.5188$ |
| Figure S1F | Two-way ANOVA | $F(1, 16) = 0.8662, P = 0.3658$ |
| Figure S1G | Two-tailed Student's t test | $t = 0.2243, P = 0.8254$ |
| Figure S1H | Two-tailed Student's t test | $t = 0.3455, P = 0.7342$ |
| Figure S1I | Two-tailed Student's t test | $t = 0.3223, P = 0.7514$ |
| Figure S1J | Two-way ANOVA | $F(1, 14) = 2.668, P = 0.1247$ |
| Figure S1K | Two-tailed Student's t test | $t = 2.464, P = 0.0240$ |
| Figure S1L | Two-tailed Student's t test | $t = 0.2667, P = 0.7928$ |
| Figure S1M | Two-tailed Student's t test | $t = 0.2226, P = 0.8264$ |
| Figure S1N | Two-tailed Student's t test | $t = 6.705, P < 0.0001$ |
| Figure 2D | Two-tailed Student's t test | $t = 3.107, P = 0.0036$ |
| Figure 2E | Two-way ANOVA | $F(1, 38) = 31.89, P < 0.0001$ |
| Figure 2G | Two-tailed Student's t test | $t = 0.9918, P = 0.3292$ |
| Figure 2H | Two-way ANOVA | $F(1, 30) = 7.577, P = 0.0099$ |
| Figure 2K | Two-tailed Student's t test | $t = 1.854, P = 0.0729$ |
| Figure 2L | Two-way ANOVA | $F(1, 27) = 18.22, P = 0.0002$ |
| Figure 3A<br>middle | Two-tailed Student's t test | $t = 5.166, P < 0.0001$ |
| Figure 3A<br>right | Two-tailed Student's t test | $t = 2.490, P = 0.0201$ |
| Figure 3B<br>middle | Two-tailed Student's t test | $t = 2.997, P = 0.0059$ |
| Figure 3B<br>right | Two-tailed Student's t test | $t = 1.191, P = 0.2446$ |
| Figure 3D | Two-tailed Student's t test | $t = 3.422, P = 0.0091$ |
| Figure 3F | Two-tailed Student's t test | $t = 2.324, P = 0.0452$ |
| Figure 3H | Two-tailed Student's t test | $t = 2.701, P = 0.0088$ |
| Figure 3I | Two-tailed Student's t test | $t = 0.2994, P = 0.7656$ |
| Figure 3K | Two-tailed Student's t test | $t = 2.224, P = 0.0292$ |
| Figure 3L | Two-tailed Student's t test | $t = 1.268, P = 0.2088$ |
| Figure 3M | One-way ANOVA followed by<br>Tukey's post hoc test | $F = 11.79, P < 0.0001$<br>KO vs. WT: $P = 0.0066$<br>KO + U0126 vs. KO: $P = 0.0009$ |
| Figure 3N | One-way ANOVA followed by<br>Tukey's post hoc test | $F = 5.654, P = 0.0021$<br>Nivo vs. IgG: $P = 0.0075$<br>Nivo + U0126 vs. Nivo: $P = 0.0100$ |
| Figure S3A | Two-tailed Student's t test | $t = 0.05485, P = 0.9580$ |
| Figure S3B | Two-tailed Student's t test | $t = 1.552, P = 0.1717$ |
| Figure S3C | Two-tailed Student's t test | $t = 6.849, P < 0.0001$ |
| Figure 4D | Two-way ANOVA | $F(1, 35) = 20.25, P < 0.0001$ |
| Figure 5B | One-way ANOVA followed by<br>Tukey's post hoc test | $F = 25.70, P < 0.0001$<br>cKO-neuron vs. control: $P = 0.0004$<br>cKO-microglia vs. cKO-neuron:<br>$P < 0.0001$ |
| Figure 5C | Two-tailed Student's t test | $t = 4.054, P = 0.0016$ |
| Figure 5D | Two-way ANOVA | $F(1, 13) = 16.73, P = 0.0013$ |
| Figure 5E | Mann-Whitney test | $U = 10, P = 0.0401$ |
| Figure 5F | Mann-Whitney test | $U = 13.5, P = 0.0892$ |
| Figure 5G | Two-tailed Student's t test | $t = 0.7668, P = 0.4569$ |
| Figure 5I | Two-tailed Student's t test | $t = 6.183, P = 0.0002$ |
| Figure 5J | Two-tailed Student's t test | $t = 4.878, P = 0.0009$ |
| Figure 5L | One-way ANOVA followed by Tukey's post hoc test | $F = 9.573, P = 0.0024$<br>cKO-microglia vs. control: $P = 0.0047$<br>cKO-neuron vs. cKO-microglia: $P = 0.0067$ |
| Figure 5M | Two-tailed Student's t test | $t = 0.8823, P = 0.3886$ |
| Figure 5N | Two-way ANOVA | $F(1, 19) = 0.1186, P = 0.2898$ |
| Figure 5O | Mann-Whitney test | $U = 40.5, P = 0.3225$ |
| Figure 5P | Mann-Whitney test | $U = 38.5, P = 0.2503$ |
| Figure 5Q | Two-tailed Student's t test | $t = 0.9261, P = 0.3660$ |
| Figure S5D | Two-tailed Student's t test | $t = 3.945, P = 0.0009$ |
| Figure S5E | Two-way ANOVA | $F(1, 20) = 18.99, P = 0.0003$ |
| Figure S5F | Mann-Whitney test | $U = 12, P = 0.0008$ |
| Figure S5G | Mann-Whitney test | $U = 12, P = 0.0005$ |
| Figure S5H | Two-tailed Student's t test | $t = 1.284, P = 0.2139$ |
| Figure S5K | Two-way ANOVA | $F(1, 13) = 1.235, P = 0.2865$ |
| Figure S5L | Two-tailed Student's t test | $t = 1.142, P = 0.2739$ |
| Figure S5M | Two-tailed Student's t test | $t = 0.4004, P = 0.6953$ |
| Figure S5N | Two-tailed Student's t test | $t = 0.8518, P = 0.4097$ |
| Figure S5O | Two-tailed Student's t test | $t = 0.5880, P = 0.5674$ |
| Figure S5R | Two-way ANOVA | $F(1, 12) = 0.7189, P = 0.4131$ |
| Figure S5S | Two-tailed Student's t test | $t = 1.486, P = 0.1630$ |
| Figure S5T | Two-tailed Student's t test | $t = 1.982, P = 0.0708$ |
| Figure S5U | Two-tailed Student's t test | $t = 0.5825, P = 0.5710$ |
| Figure S5V | Two-tailed Student's t test | $t = 0.3078, P = 0.7635$ |
| Figure 6E | Two-tailed Student's t test | $t = 8.906, P < 0.0001$ |
| Figure 6F | Two-way ANOVA | $F(1, 21) = 14.89, P = 0.0009$ |
| Figure 6G | Mann-Whitney test | $U = 20, P = 0.0035$ |
| Figure 6H | Mann-Whitney test | $U = 27.5, P = 0.0134$ |
| Figure 6I | Two-tailed Student's t test | $t = 1.133, P = 0.2710$ |
| Figure 6K | Two-tailed Student's t test | $t = 4.191, P = 0.0019$ |
| Figure 6L | Two-tailed Student's t test | $t = 4.963, P = 0.0006$ |
| Figure S7C<br>upper | Two-tailed Student's t test | $t = 4.963, P = 0.4293$ |
| Figure S7C<br>bottom | Two-tailed Student's t test | $t = 4.963, P = 0.7147$ |
| Figure S7D | Two-way ANOVA | $F(1, 21) = 0.02744, P = 0.8700$ |
| Figure S7E | Two-tailed Student's t test | $t = 0.4572, P = 0.6522$ |
| Figure S7F | Two-tailed Student's t test | $t = 1.167, P = 0.2562$ |
| Figure S7G | Two-tailed Student's t test | $t = 0.3724, P = 0.7133$ |
| Figure S7H | Two-tailed Student's t test | $t = 0.7982, P = 0.4337$ |
| Figure 7B | One-way ANOVA followed by<br>Tukey's post hoc test | $F = 29.62, P < 0.0001$<br>KO Sham vs. WT sham: $P = 0.0005$<br>KO TBI vs. WT TBI: $P < 0.0001$<br>WT TBI vs. WT sham: $P = 0.0003$<br>KO TBI vs. KO Sham: $P = 0.0050$ |
| Figure 7C | Two-way ANOVA | KO Sham vs. WT sham: $P < 0.0001$<br>KO TBI vs. WT TBI: $P = 0.0112$<br>WT TBI vs. WT sham: $P = 0.0004$<br>KO TBI vs. KO Sham: $P = 0.0004$ |
| Figure 7D | Kruskal-Wallis test followed by<br>Dunn's post hoc test | Kruskal-Wallis statistic = 18.29, $P = 0.0004$<br>KO TBI vs. WT TBI: $P = 0.0424$ |
| Figure 7E | Kruskal-Wallis test followed by<br>Dunn's post hoc test | Kruskal-Wallis statistic = 12.05, $P = 0.0072$<br>KO TBI vs. WT TBI: $P = 0.0402$ |
| Figure 7F | One-way ANOVA followed by<br>Tukey's post hoc test | $F = 4.316, P = 0.0127$<br>KO TBI vs. KO Sham: $P = 0.0131$ |
| Figure 7H | Two-tailed Student's t test | $t = 7.781, P < 0.0001$ |
| Figure 7I | Two-way ANOVA | $F(1, 14) = 9.623, P = 0.0078$ |
| Figure 7J | Mann-Whitney test | $U = 13, P = 0.0468$ |
| Figure 7K | Mann-Whitney test | $U = 21.5, P = 0.3002$ |
| Figure 7L | Two-tailed Student's t test | $t = 0.06360, P = 0.9502$ |
| Figure 7M | Two-tailed Student's t test | $t = 1.394, P = 0.1851$ |
| Figure 7N | Two-tailed Student's t test | $t = 0.9678, P = 0.3495$ |
| Figure 7O | Two-tailed Student's t test | $t = 0.1476, P = 0.8848$ |
| Figure 7P | Two-tailed Student's t test | $t = 0.9696, P = 0.3487$ |
| Figure 7Q | Two-tailed Student's t test | $t = 0.8938, P = 0.3866$ |
| Figure 7R | Two-tailed Student's t test | $t = 1.292, P = 0.2172$ |
| Figure S8B | Two-tailed Student's t test | $t = 1.576, P = 0.1461$ |
| Figure S8D | Two-tailed Student's t test | $t = 1.590, P = 0.1341$ |
| Figure S8E | Two-tailed Student's t test | $t = 2.054, P = 0.0591$ |
| Figure S8F | Two-tailed Student's t test | $t = 0.1507, P = 0.8824$ |
| Figure S8G | Two-tailed Student's t test | $t = 1.160, P = 0.2656$ |
| Figure S8H | Two-tailed Student's t test | $t = 0.9762, P = 0.3455$ |

